# Mitochondrial malate metabolism acts as a control hub for photosynthesis and carbon-nitrogen balance in Arabidopsis

**DOI:** 10.64898/2026.03.02.709023

**Authors:** María del Pilar Martinez, Ioana Nica, Ke Zheng, Noah Ditz, Julie A. V. S. de Oliveira, Pedro Barreto, Nina Blum, Philipp Westhoff, Boas Pucker, Holger Eubel, Iris Finkemeier, Markus Schwarzländer, Veronica G. Maurino

## Abstract

Malate is a central metabolite in plant energy metabolism and biosynthesis and serves as a major carrier of carbon and reducing equivalents between chloroplasts, the cytosol, and mitochondria. However, how individual malate-converting systems contribute to physiology in specific subcellular compartments remains incompletely understood. Here, we investigated the impact of combined loss of mitochondrial malate dehydrogenase (MDH) and NAD-dependent malic enzyme (NAD-ME) activity in *Arabidopsis thaliana* by integrating reverse genetics, physiological analyses, transcriptomics, quantitative proteomics, and metabolite profiling. Specifically, we generated triple mutants (*mdh1xme1xme2*) lacking the predominant mitochondrial isoform MDH1 together with both NAD-ME subunits, thereby reducing overall mitochondrial malate conversion capacity. By growing plants under contrasting photoperiod and irradiance regimes to vary photosynthetic demand on malate-linked fluxes, we uncovered a conditional phenotype that was most pronounced under short-day/low-light conditions. Under these conditions, *mdh1xme1xme2* exhibited impaired growth and photosynthetic performance, accompanied by cytosolic redox imbalance and altered chloroplast ultrastructure. Transcriptomic profiling revealed that low light unmasks a dawn-phase bottleneck in establishing photosynthetic and redox homeostasis. Consistent with this, the low-light plastid proteome revealed a reallocation away from chloroplast translation and photosynthetic capacity toward proteome maintenance, photoprotection/repair, and iron/ROS management, consistent with a protective acclimation state that nevertheless constrains carbon gain under energy limitation. Low light also triggered C/N imbalance and ammonium accumulation in the mutants. In contrast, increasing irradiance or extending the photoperiod largely alleviated these defects. Together, our results identified mitochondrial malate conversion capacity as a key control point coupling respiratory energy supply and redox homeostasis to photosynthetic metabolism when photosynthetic energy input is limiting.

## Introduction

Malate is a central intermediate of the tricarboxylic acid (TCA) cycle in plant cells, with functions that extend far beyond its role in mitochondrial energy metabolism (Maurino and Engqvist, 2015). It participates in a wide range of essential processes, including photosynthesis, fatty acid oxidation, stomatal regulation, nitrogen fixation, amino acid biosynthesis, ion balance, phosphorus, iron uptake, and aluminum tolerance (Martinoia and Rentsch, 1994; Kochian, 1995; Maurino and Engqvist, 2015; Drincovich et al., 2016). Within mitochondria, malate is a direct intermediate of the TCA cycle, and its oxidation generates NAD(P)H that supports ATP production required for growth and development (Møller and Rasmusson, 1998). Beyond this canonical function, malate also participates in redox-coupled reactions that connect carbon and nitrogen metabolism – for example, its decarboxylation to pyruvate by NAD-malic enzyme (NAD-ME) and its reversible interconversion with oxaloacetate (OAA) by malate dehydrogenase (MDH) (Scheibe, 2004).

Maintaining redox balance across subcellular compartments is essential for the proper functioning of cofactor-dependent reactions that coordinate metabolism and optimize photosynthesis. While NAD(P)^+^ transporters have been identified in chloroplasts, mitochondria, and peroxisomes, their low affinity for NAD(P)H restricts the direct transfer of reducing equivalents (Palmieri et al., 2009). Instead, organelles predominantly rely on malate transport as an indirect mechanism to shuttle reducing power. In a process known as malate valve, malate and OAA are exchanged between cell compartments via specific transmembrane translocators, and the two metabolites can be interconverted at either side of the membrane by MDH activity (Selinski and Scheibe, 2019). Because the reaction is fully reversible, the net flux direction is governed by mass action, that is, by metabolite concentrations shaped by cellular demand. By facilitating these redox exchanges, malate valves are thought to maintain metabolic balance between compartments and, importantly, support optimal photosynthetic performance (Raghavendra and Padmasree, 2003).

The reversible interconversion of malate and OAA is catalyzed by NAD(P)-MDHs (Gietl, 1992). The *A. thaliana* Columbia-0 genome encodes nine NAD(P)-MDHs distributed across different organelles and the cytosol. In Arabidopsis mitochondria, two nuclear encoded NAD-MDH are localized, *MDH1* (At1g53240) and *MDH2* (At3g15020), with MDH1 being the predominant MDH enzyme (Lee et al., 2008; Tomaz et al., 2010; Hüdig et al., 2015; Fuchs et al., 2020). MDH functions as a homodimer and is regulated by adenine nucleotides as well as acetylation (Yoshida and Hisabori, 2016; Balparda et al., 2022; Zheng et al., 2025). In illuminated mature leaves, MDH is thought to operate mainly in the direction of OAA reduction and NADH oxidation to export malate from the mitochondrial matrix to the cytosol (Shameer et al., 2019). At the same time, the TCA cycle is partially inactive due to inhibition of mitochondrial pyruvate dehydrogenase (PDH) complex, promoting mitochondrial OAA import and malate efflux that support redox coupling between the matrix and other subcellular compartments (Thum et al., 2004; Rasmusson and Escobar, 2007; de Oliveira Dal’Molin et al., 2010; Bykova et al., 2014; Hüdig et al., 2015). At night, when photosynthesis ceases and the demand for ATP increases, the TCA cycle resumes cyclic operation, with MDH primarily catalyzing malate oxidation coupled to NAD^+^ reduction (Sweetlove et al., 2010). During this period, reducing equivalent exchange between chloroplast and mitochondrial via malate shuttles is negligible due to the cessation of photosynthesis (Kromer, 1995; Raghavendra and Padmasree, 2003; Scheibe, 2004; Selinski and Scheibe, 2019; Zhao et al., 2020). This concept is further supported by recent evidence of a shift to a more reduced cyto-nuclear NAD pool in loss-of-function lines of mitochondrial MDH1 or MDH2 during darkness (Zheng et al., 2025). A recent succession of reports have put forward an appealing working model of ‘malate circulation’ that includes the opposite direction of flux, i.e. OAA import into the mitochondria in the light in Arabidopsis and the C4 species maize (Wu et al., 2015; Zhao et al., 2018; Zhao et al., 2020; Jiang et al., 2023). Whether such a flux mode can operate in *Arabidopsis* leaves in vivo under specific conditions remains to be established and will require careful consideration of cellular metabolic constraints.

In addition to MDH, NAD-ME metabolizes malate in mitochondria by catalyzing its NAD^+^-dependent decarboxylation to pyruvate (Tronconi et al., 2008). NAD-ME is an accessory enzyme of the TCA cycle by enabling conversion of malate to pyruvate and functions primarily as a heterodimer composed of ME1 (At2g13560) and ME2 (At4g00570) (Tronconi et al., 2010). Flux prediction models of the TCA cycle suggest that a stable metabolic state can be maintained through a cyclic flux in which malate, rather than pyruvate, serves as the primary substrate (Steuer et al., 2007). In this scenario, NAD-ME would convert malate to pyruvate, thereby feeding the cycle through the PDH complex. However, empirical evidence suggests that NAD-ME contributes minimally to the pyruvate pool when the mitochondrial pyruvate carrier (MPC) supplies pyruvate to the TCA cycle (Dieuaide-Noubhani et al., 1995; Williams et al., 2008; Le et al., 2022). NAD-ME-derived pyruvate is preferentially redirected to the cytosol potentially contributing to acetyl-CoA formation in plastids for fatty acid biosynthesis (Camp and Randall, 1985; Post-Beittenmiller et al., 1992; Ke et al., 2000; Le et al., 2022). Moreover, pyruvate can be converted to phosphoenolpyruvate (PEP) in the cytosol and plastids to maintain pH balance under water stress (Netting, 2000; Santelia and Lawson, 2016).

Taken together, MDH and NAD-ME provide mitochondria with substantial flexibility to modulate carbon fluxes across the day-night cycle. Analyses of individual loss-of-function mutants of MDH (*mdh1* and *mdh2*) and NAD-ME (*me1* and *me2*), as well as *me1*x*me2* double mutants, revealed metabolic deviations from the wild type but no apparent effects on plant growth (Tronconi et al., 2008; Tomaz et al., 2010). In contrast, *mdh1*x*mdh2* double mutants show reduced growth under both long- and short-day conditions, accompanied by decreased net CO_2_ assimilation and increased leaf respiration rates, effects that occur independently of photoperiod (Tomaz et al., 2010). Despite these deep insights and advanced working models, the physiological importance of mitochondrial malate oxidation and the potential consequences of its impairment are not immediately evident due to the flux-network-nature of central metabolism. Moreover, recent conflicting models demand dedicated experimental dissection of this major hub of central plant metabolism.

Here, we investigated the impact of combined MDH and NAD-ME loss-of-function in *A. thaliana* by integrating reverse genetics, physiological analyses, transcriptomics, quantitative proteomics, and metabolite profiling. Specifically, we generated triple mutants (*mdh1xme1xme2*) lacking the predominant mitochondrial isoform MDH1 together with both NAD-ME subunits to diminish overall mitochondrial malate conversion capacity. Growth under different loads of photosynthetic metabolism as established through different light regimes revealed a short-day/low-light specific phenotype, characterized by cytosolic redox imbalance, altered chloroplast ultrastructure, and impaired photosynthetic performance and CO_2_ assimilation.

## Results and Discussion

### Disruption of mitochondrial malate utilization by combining *MDH1* and *NAD-ME* loss of function

To investigate how impaired mitochondrial malate utilization affects plant performance, we targeted the predominant mitochondrial MDH1 along with the subunits of NAD-ME, ME1 and ME2. Previously reported T-DNA insertion lines for *MDH1* (*mdh1.1*; (Tomaz et al., 2010)), hereafter *mdh1*), *ME1* (*me1*; (Tronconi et al., 2008)), and *ME2* (*me2.1* and *me2.2*; (Tronconi et al., 2008)) were re-validated to map insertion sites and assayed for transcript abundance by RT-qPCR (Fig. 1A and Suppl. Table 1). All lines are confirmed as knockouts at the transcript level except for *me2.2*, which retains 20% of residual amounts of *ME2* transcript (Fig. 1B). We hence generated two distinct triple loss-of-function lines, *mdh1xme1xme2.1* and *mdh1xme1xme2.2,* differing in the *me2* allele, by genetic crosses (Suppl. Table 1). Attempts to produce a quadruple mutant with an additional T-DNA insertion in *MDH2* (*mdh2*; (Tomaz et al., 2010)) were unsuccessful despite our repeated efforts, suggesting that additional more severe disruption of mitochondrial malate conversion capacity is detrimental. Severe defects in early development have previously been established when MDH1 and MDH2 are both disrupted (Sew et al., 2016).

**Figure 1.**
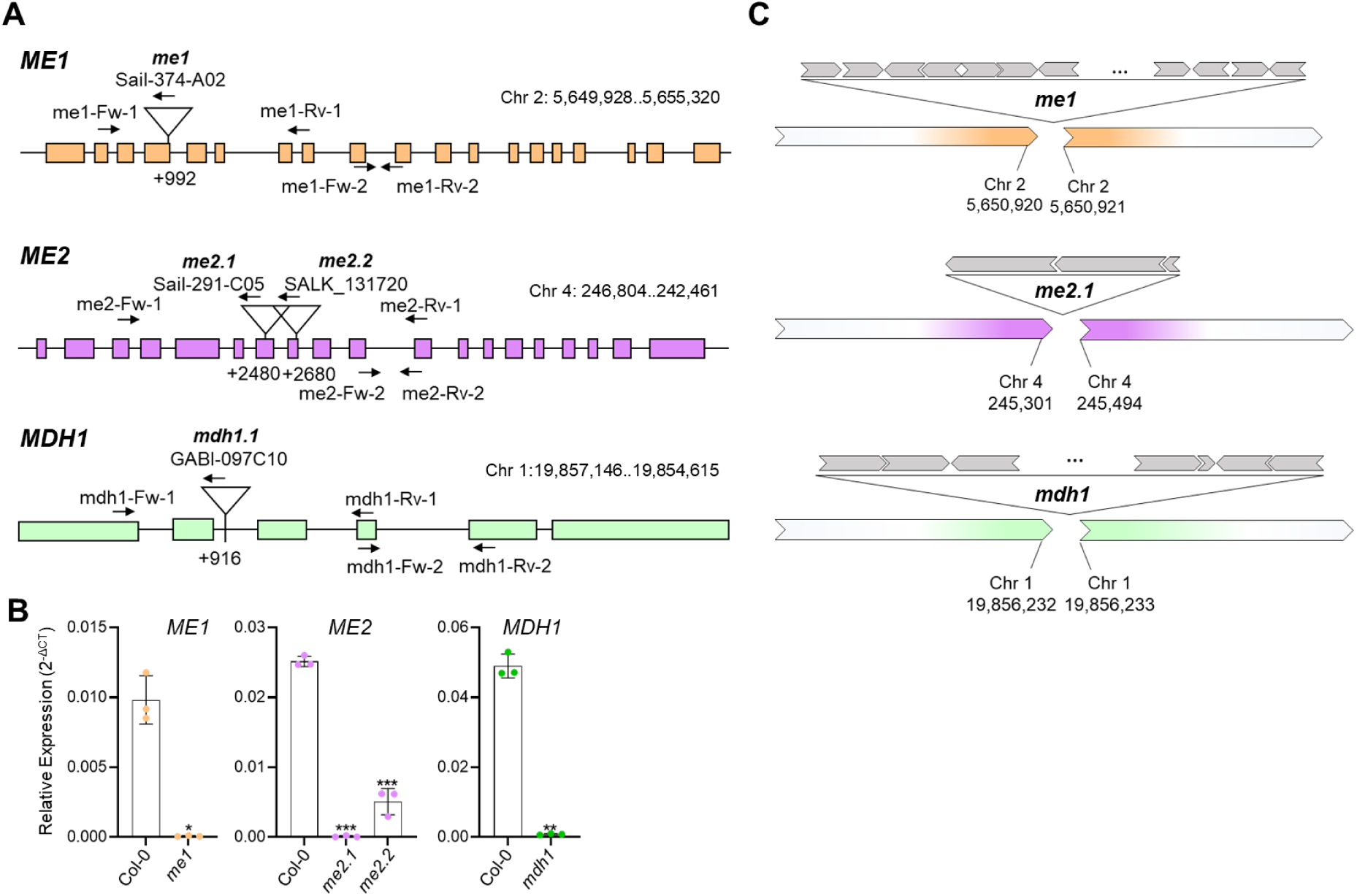
T-DNA insertion lines of *A. thaliana* mitochondrial *NAD-ME* and *MDH1*. **A)** Gene models of *ME1*, *ME2*, and *MDH1* indicating the confirmed T-DNA insertion sites. Locations of primer binding sites used for zygosity analysis and RT-qPCR (Suppl. Table 2) are indicated by black arrows. Rectangles represent exons and spaces between them introns. **B)** Relative expression levels of *ME1*, *ME2*, and *MDH1* in rosette leaves of 40-day-old Col-0 and single insertional mutants grown in SD at LL, harvested 1 h before onset of the light. Data are means ± SD (n = 3). Significance versus Col-0 was assessed by Welch’s t-test (p < 0.05, *; p < 0.01, **; p < 0.001, ***; p < 0.0001, ****). **C)** Chromosomal positions and number of T-DNA insertions in each candidate gene as determinate by ONT sequencing. T-DNA insertions are represented in gray and their orientation is indicated by arrows. The length of the segments is not scaled.

We sequenced the *mdh1xme1xme2.1* genome using long-read whole-genome sequencing by Oxford Nanopore Technology (ONT). T-DNA arrays mapped to the expected loci - At1g53240 (*MDH1*), At4g00570 (*ME2*), and At2g13560 (*ME1*) (Fig. 1C). In addition, *mdh1xme1xme2.1* carries a T-DNA insertion within the 3’-UTR of At5g62090 (*SLK2*), positioned 42 bp upstream of the annotated gene end (Suppl. Fig. 1A). Despite this insertion, transcriptome analysis shows no significant differences in *SLK2* expression relative to wild type (Suppl. Fig. 1B). The triple mutant also harbors a T-DNA insertion in the transposable element AT2TE41910, which does not affect protein encoding genes. Because T-DNA mutagenesis can be accompanied by genomic rearrangements, which were reported for several GABI-Kat lines (Pucker et al., 2021), we assessed read-depth coverage to test for large-scale rearrangement in *mdh1xme1xme2.1* and found no evidence for such events (Suppl. Fig. 2).

### Transcript dynamics of *MDH1* and *ME* at light transitions

TCA cycle activity is tightly regulated by different external factors, such light intensity, through the regulation of the expression of the TCA cycle enzymes (Thum et al., 2004; Tcherkez et al., 2005; Popov et al., 2010). To assess how irradiance shapes the transcript dynamics of mitochondrial MDH and ME isoforms in Col-0 at light-dark transitions, we performed RT-qPCR of plants grown in short days (SD) at 60 (LL) or 120 (NL) µmol photons m^-2^ s^-1^. Leaf tissue was harvested 1 h before lights-on (-1), 1 h after lights-on (+1), and 7 h into the light period (+7).

In Col-0, *MDH1* transcript levels are consistently higher than *MDH2* under both light intensity regimens, with the difference being most pronounced during the day (Fig. 2A and B). Likewise, *ME2* generally exceeds *ME1*, except at +1 at NL, where *ME1* surpasses *ME2*. While expression patterns at -1 and +7 are similar between LL and NL, a pronounced divergence occurs at +1: transcripts at LL are substantially lower than at NL. These data highlight the impact of light intensity on the expression of *MDH1* and both *ME* genes.

**Figure 2.**
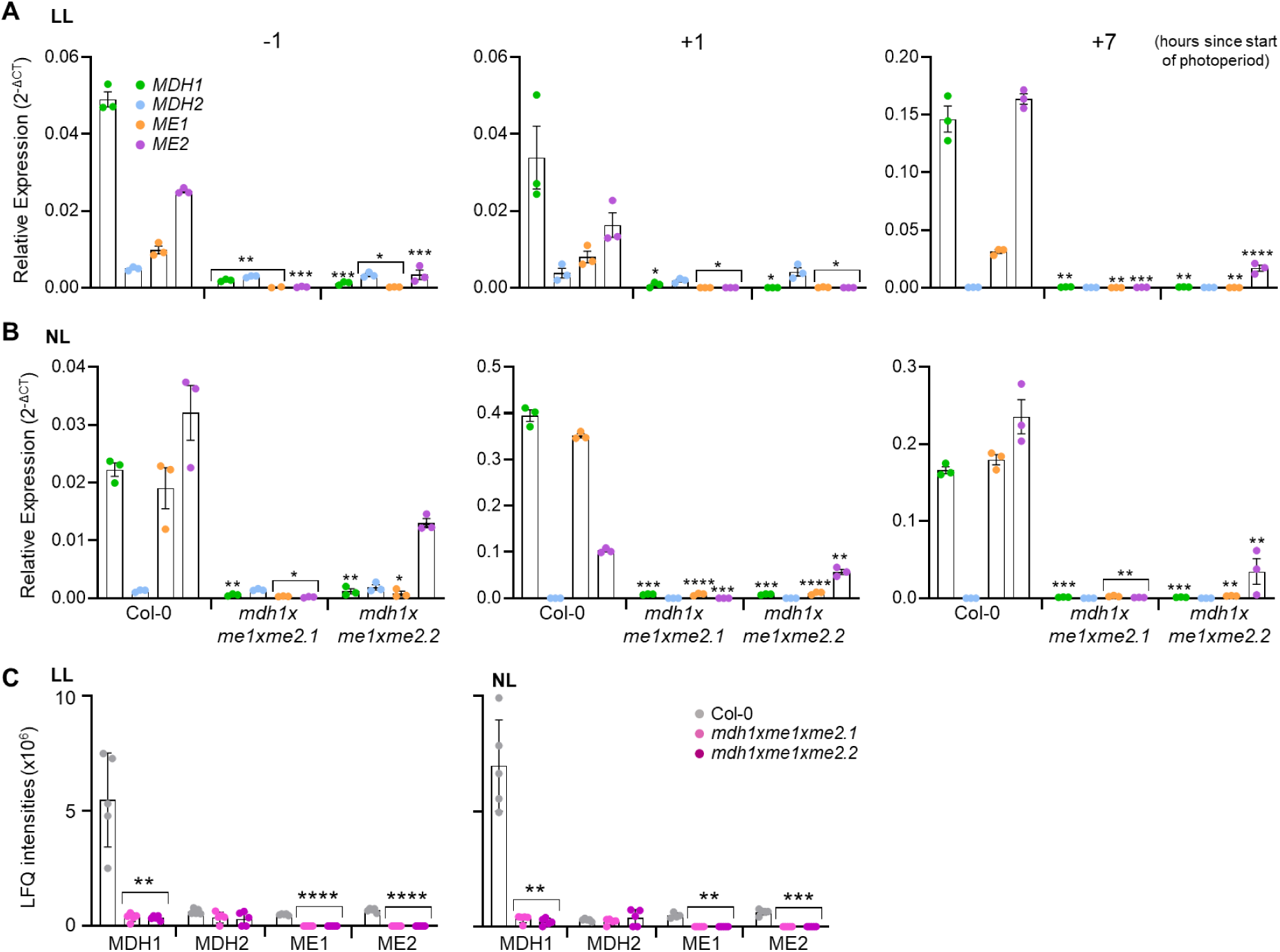
MDH and ME transcript and protein abundance under short day conditions. **A)** Relative transcript levels at low light (LL). **B)** Relative transcript levels at normal light (NL). Leaf tissue was harvested 1 hour before lights-on (-1), 1 hour after lights-on (+1), and 1 hour before end of day (+7). Data are means ± SEM (n = 3). **C)** Label-free quantification (LFQ) in plants grown in SD at LL or NL. Bars show mean ± SD (n = 5). Significance versus Col-0 was assessed by Welch’s t-test (p < 0.05, *; p < 0.01, **; p < 0.001, ***; p < 0.0001, ****). Bars under the brackets present the same significance compared with Col-0.

*MDH1* and *ME1* transcripts are depleted in both triple mutant lines (Fig. 2A and B). *ME2* transcript levels were also diminished to background levels in *mdh1xme1xme2.1*, whereas in *mdh1xme1xme2.2* residual *ME2* transcripts decreased in abundance by 50-90 % but detectable at most time points, consistent with the *me2.2* knockdown allele.

### Lack of MDH1 and ME-derived peptides in *mdh1xme1xme2*

Because residual ME2 transcript levels in the *me2.2* line could potentially sustain protein accumulation, we assessed protein abundance by LC-MS/MS proteomics. Multiple ME1- and ME2-derived peptides were detected in wild-type samples but none in the triple mutants (Fig. 2C). Notably, despite the residual *ME2* transcript in the *me2.2* background, no ME2-derived-peptides were detected. Similarly, MDH1-derived peptides were readily detected in wild type but no MDH1-unique peptides were found in the triple mutants. Collectively, the transcript and proteomic data indicate that the triple mutants lack detectable mitochondrial MDH1, ME1, and ME2 protein, making both lines suitable to study the impact of diminished mitochondrial malate conversion capacity.

MDH2 is the only remaining mitochondrial MDH isoform in both triple mutant lines. We therefore assessed whether MDH2 shows compensatory changes in transcript or protein abundance in the absence of MDH1 and NAD-ME. *MDH2* transcript levels were indistinguishable from wild type at all time points and under both light intensities (Fig. 2A and B), and MDH2 protein abundance likewise remained unchanged (Fig. 2C). These results indicate that MDH2 expression does not compensate for the loss of MDH1 and ME in mitochondrial malate oxidation/oxaloacetate reduction.

### Combined loss of MDH1 and NAD-ME causes a photoperiod- and irradiance-dependent growth defect in *A. thaliana*

Because subcellular malate flux is tightly coupled to photosynthetic activity (Herrmann et al., 2021), we aimed at titrating the load of assimilatory carbon metabolism by assessing plant development under different photoperiods and irradiance regimens. Under long day conditions (LD) at either LL or NL, as well as under SD at NL, the triple mutants were phenotypically indistinguishable from wild type (Fig. 3A and B). By contrast, under SD combined with LL, the triple mutants displayed pale-green coloration and reduced rosette size relative to wild type (Fig. 3A and B; Suppl. Fig. 3). All other genotypes tested showed wild-type-like growth under both LD-NL, and SD-LL conditions (Suppl. Fig 3). Consistent with the impaired vegetative growth under SD-LL, the wild-type initiated bolting at about 42 days after sowing, whereas triple mutants bolted 14 days later (Fig. 3C), indicating a marked delay in the vegetative-to-reproductive transition. This bolting delay was not observed under any other condition examined. Collectively, these results show that the combined loss of MDH1 and NAD-ME confers a developmental defect that manifests specifically under the SD-LL regime, revealing a profound link of mitochondrial malate handling with photoperiod and light intensity.

**Figure 3.**
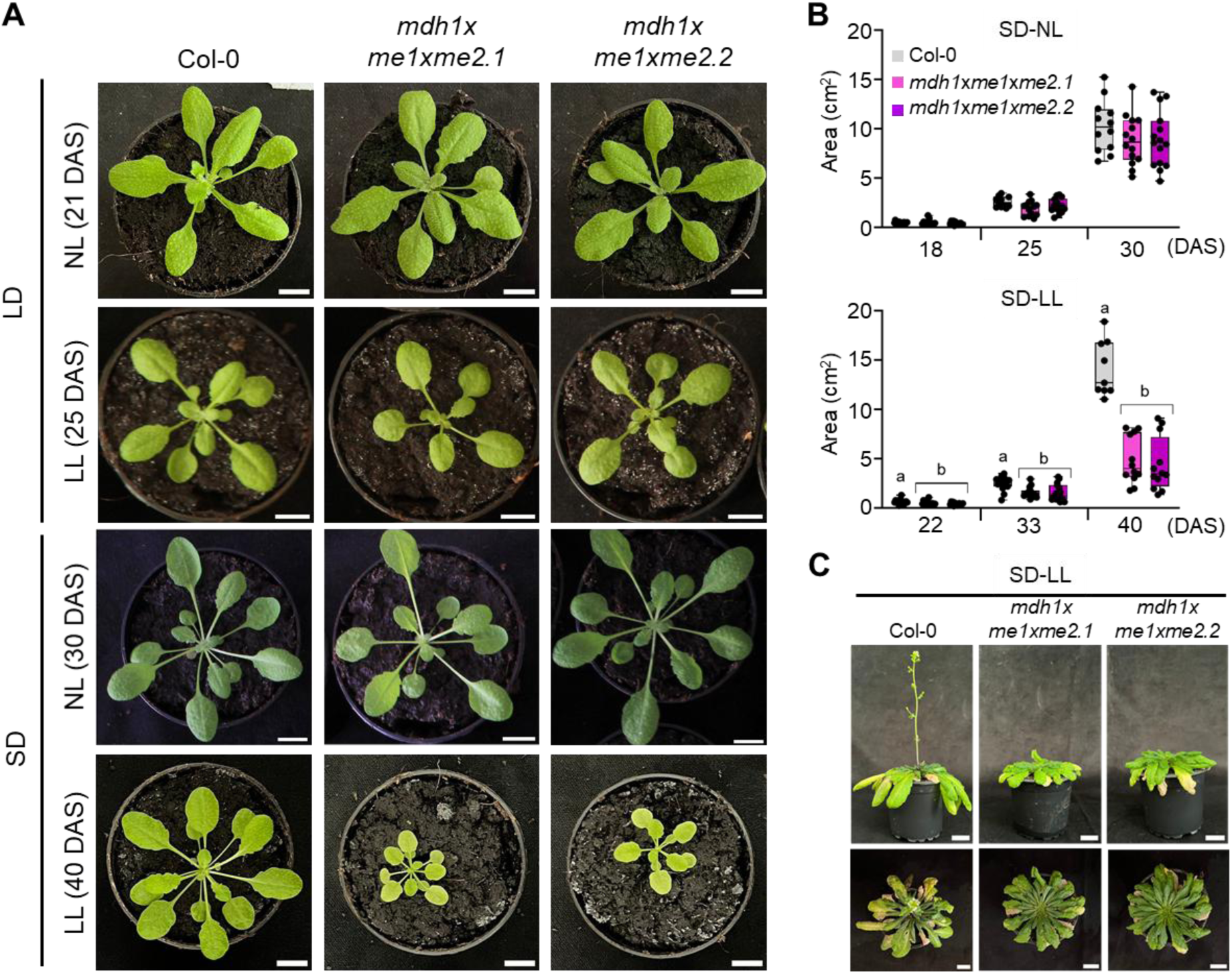
Growth of plants under different photoperiods and light intensities. **A)** Representative plants of each genotype grown during 21 days in LD at NL, 25 days in LD at LL, 30 days in SD at NL, and 40 days in SD at LL. Representative plants of additional genotypes are shown in Supplementary Figure 2. Scale bar = 1 cm. **B)** Rosette area at different days after sowing (DAS) under SD at NL and LL. Each point inside the box represents one biological replicate (n = 10). Statistics were performed separately for each day using one-way ANOVA followed by Tukey’s post-hoc test. Different letters indicate statistically significant differences (*p* < 0.05). **C)** Representative plants of each genotype at 60 DAS in SD at LL. Scale bar = 1 cm. LD: long day; LL: low light; NL: normal light; SD: short day.

### Compromised photosynthesis and altered chloroplasts ultrastructure in *mdh1*x*me1xme2*

As the mutants showed small pale green rosettes under SD-LL, we quantified key photosynthetic parameters as indicators of carbon assimilation competence. The mutants exhibited lowered photosynthetic capacity under SD-LL but performed similarly to wild type at NL (Fig. 4A and B). At LL, imaging of Fv/Fm (maximum quantum yield of PSII) revealed that the reduction in photosynthetic efficiency was primarily restricted to mesophyll tissue of older leaves, whereas vasculature bundles largely retained their photosynthetic competence (Fig. 4A). Consistently, the mutants showed lower PSII quantum yield (ΦPSII) and electron transport rate (ETR) compared to wild type, demonstrating impaired PSII function and reduced photochemical efficiency. By contrast, under NL these parameters were indistinguishably from wild type. Non-photochemical quenching (NPQ) is elevated in the triple mutants at LL, indicating that they are able to efficiently capture light energy, but present problems converting it to chemical energy in the reaction center. Contrary, NPQ values remained at wild type levels at NL (Fig. 4B).

**Figure 4.**
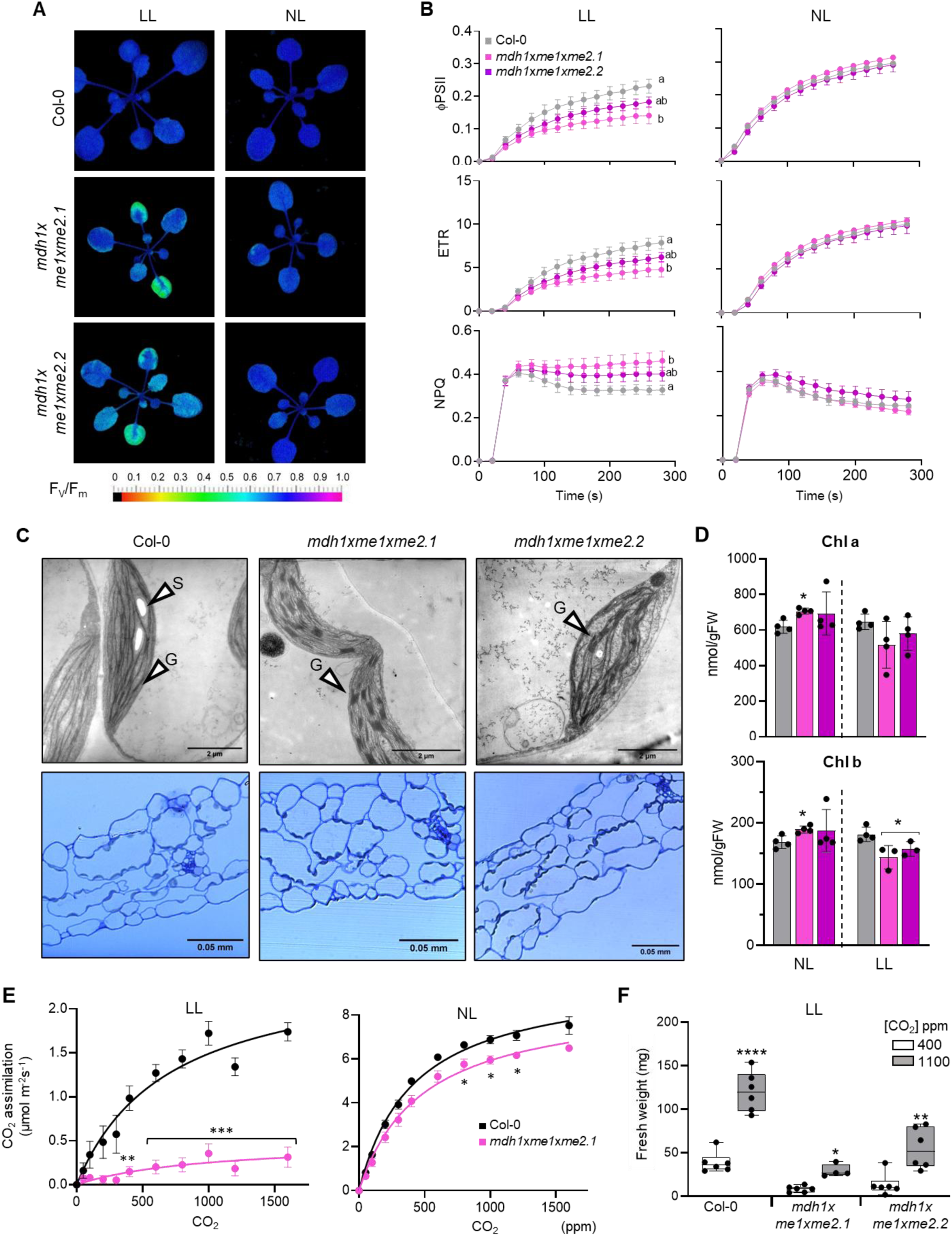
Chloroplast functionality and morphology from plants grown in short days at low or normal light. **A)** Representative false-color images of maximum quantum yield of PSII (Fv/Fm) in 5-week-old plants. **B)** Photosynthetic parameters derived from chlorophyll fluorescence imaging in A: ΦPSII (operational quantum yield of PSII), ETR (electron transport rate), and NPQ (non-photochemical quenching). Values are means ± SEM (n = 3). **C)** Transmission electron micrographs of leaf sections from 40-day-old plants grown in SD at LL and harvested at the end of the night. G, grana; S, starch granule. Bottom panels: semi-thin sections stained with Azur II and methylene blue to enhanced chloroplasts contrast. **D)** Chlorophyll a and b contents are shown as means ± SD from three to four biological replicates. Letters indicate significant differences (*p* < 0.05) according to one-way ANOVA followed by Tukey’s post-hoc test. Chl, chlorophyll. **E)** CO_2_ assimilation rates measured with the LI-COR 6800 system across a CO_2_ range of 0-1700 ppm at LL and NL. Values are means ± SE (n = 3-5). Statistical analysis was done using Welch’s t-test (*p* < 0.05, *; *p* < 0.01, **; *p* < 0.001, ***) F) Rosette fresh weight of Col-0 and triple mutants grown under LL SD conditions at 400 ppm CO_2_ (normal) or 1100 ppm CO_2_ (high). Values are means ± SE (n = 5-7). Testing for significant differences was performed by the Welch’s t-test. The asterisk (*) indicates significant differences between the values under normal and high CO_2_ conditions (*p* < 0.05, *; *p* < 0.01, **; *p* < 0.001, ***; p< 0.0001, ****). LL: low light; NL: normal light; SD: short day.

Ultrastructural analysis of mesophyll chloroplasts revealed increased grana stacking in *mdh1xme1xme2* relative to the wild type under LL (Fig. 4C), consistent with a potential compensatory adjustment that expands thylakoid membrane surface area and enhances photon capture efficiency. Under LL, *mdh1xme1xme2.1* plant displayed lower chlorophyll *b* content, consistent with impaired light-harvesting efficiency and the pale/yellow leaf phenotype (Fig. 4D). In contrast, under NL, both chlorophyll *a* and *b* accumulated to higher levels than in wild type, suggesting enhanced pigment biosynthesis or stability under these conditions (Fig. 4D).

### Elevated CO_2_ leads to lower CO_2_ assimilation and partial rescue of the growth defect in mdh1xme1xme2

Consistent with the growth phenotype, *mdh1xme1xme2.1* grown at LL exhibited much lower net CO_2_ assimilation than wild type, whereas at NL assimilation was closer to wild type levels (Fig. 4E).

Given that the triple mutants exhibited slower growth and reduced CO_2_ assimilation under SD-LL, yet largely recovered at higher irradiance, we next asked whether increasing CO_2_ availability could mitigate the LL growth phenotype. For that purpose, plants were grown under LL at elevated CO_2_ (1,100 ppm). Under these conditions, rosette biomass increased in both Col-0 (3-times) and the mutants (about 4-times) (Fig. 4F and Suppl. Fig 4). The higher increase in the mutant suggests greater sensitivity to CO_2_ availability than Col-0 under the cultivation conditions used here. The stronger relative response of the mutants indicates that their growth is more strongly constrained by CO_2_ availability than that of Col-0 under SD-LL. Nevertheless, elevated CO_2_ did not fully restore mutant biomass to wild-type levels (Fig. 4F and Suppl. Fig 4). Together, these data show that the triple mutants remain capable of translating increased CO_2_ supply into biomass, but that CO_2_ enrichment alone is insufficient to fully compensate for the underlying metabolic limitation.

### Dark respiration is lowered and cytosolic NAD redox state is shifted in *mdh1xme1xme2*

Since the growth defects of the mutants emerged under long nights and low irradiance, we hypothesized that mitochondrial energy supply in darkness becomes limiting under these conditions. Accordingly, we measured leaf dark respiration of plants grown under SD and found that at NL, wild type and *mdh1xme1xme2.1* exhibited similar oxygen consumption rate, whereas at LL the mutants exhibited a lower dark-respiration rate compared with the wild type (Fig. 5A and B).

**Figure 5.**
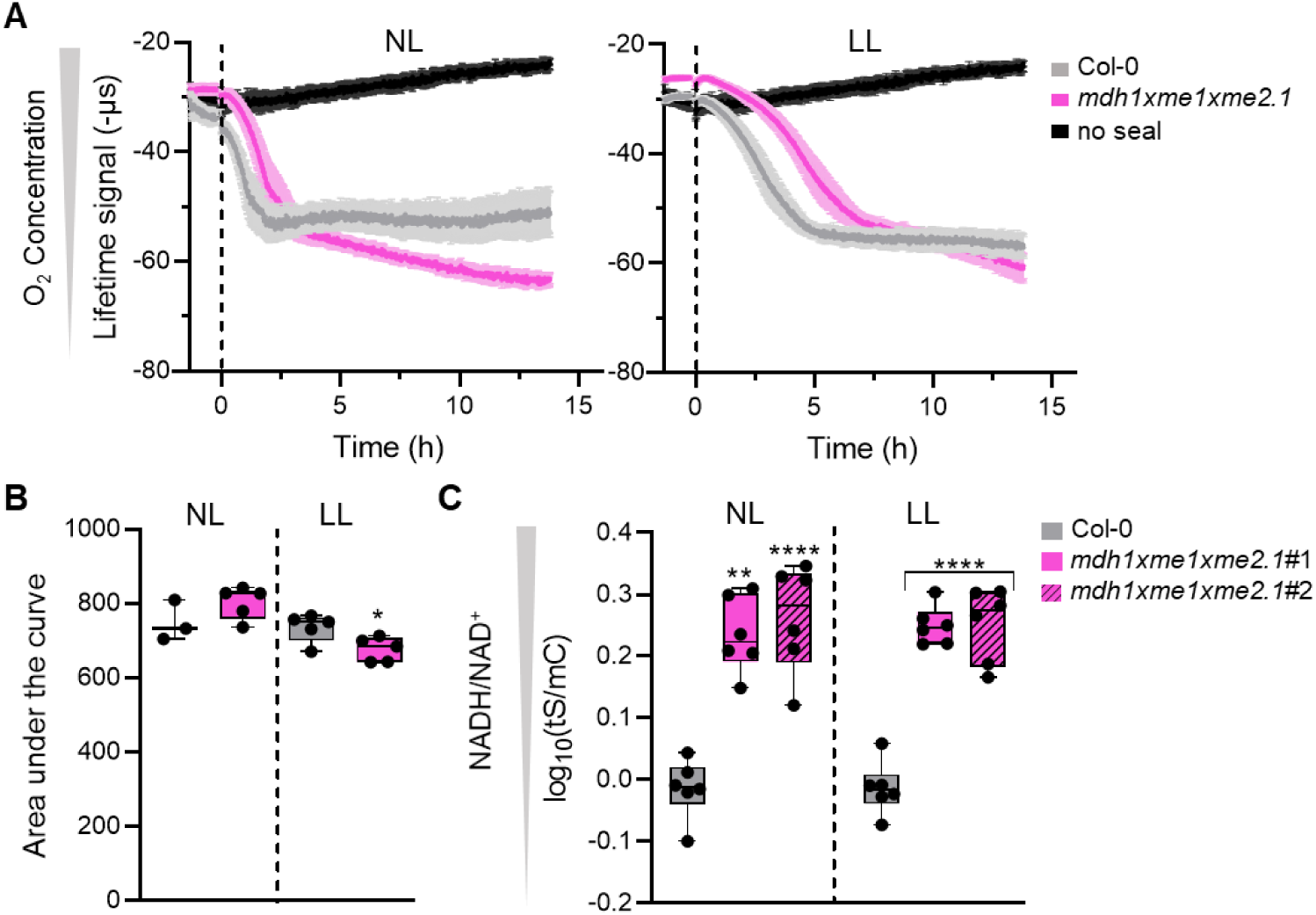
O_2_ consumption and NADH/NAD^+^ dynamics in short day conditions. **A)** Fluorescent lifetime measurements using the MitoXpress probe wells containing leaf discs from plants grown under SD at normal light (NL) and low light (LL). Data represent the mean ± SE (n = 3-5). The dashed line marks the points at which the system was sealed after reaching equilibrium. **B)** Total oxygen consumption was derived from the area under the curve using inverted curves from (A). Data are presented as the mean ± SD (n = 3-5). Statistically significant was assessed using Welch’s t-test. Asterisks denote significant differences from Col-0 (*p <* 0.05*, *; p <* 0.01*, **; p <* 0.001*, ***; p<* 0.0001*, \*\*\*\**). **C)** Cytosolic NAD redox status measured with the Peredox-mCherry (tS: tSapphire; mC: mCherry) sensor in leaf mesophyll of plants grown under SD at LL (40 DAS) or NL (30 DAS). Measurements were performed during the day in leaves after dark adaption for 2h. Data are log10-transformed, averaged ± SD (n = 6) for two independent *mdh1xme1xme2.1* presented the sensor in the cytosol lines (#1, #2). Asterisks indicate significant differences from Col-0 (*p <* 0.05*, *; p <* 0.01*, **; p <* 0.001*, ***; p<* 0.0001*, \*\*\*\**) according to the Welch’s t-test.

Given the impairment of mitochondrial malate metabolism which is intimately linked to NAD redox status, we next examined organelle-cytosol redox coupling. Col-0 and *mdh1xme1xme2.1* were transformed with the NADH/NAD⁺ biosensor Peredox-mCherry localized to the cytosol (Steinbeck et al., 2020). Ratiometric confocal imaging of leaf mesophyll tissue from plants cultivated under SD revealed a markedly more reduced cytosolic NAD pool (elevated NADH/NAD⁺) in the mutant lines relative to wild type, regardless of intensity of cultivation light (measurements were performed in leaves after dark adaption) (Fig. 5C). These results indicate a shift of the cytosolic NAD redox balance towards reduction in the mutants, consistent with altered redox shuttling between mitochondria, chloroplasts, and the cytosol. Notably, because the cytosolic NAD redox shift occurs at both LL and NL, whereas the respiration defect is specific to LL, redox status alone cannot explain the low-light growth phenotype, pointing to additional constraints that become limiting under low irradiance. High cytosolic NADH/NAD^+^ ratio was also observed in Arabidopsis *mdh1* mutants (Zheng et al., 2025) and in *N. tabacum* plants lacking mitochondrial MDH and cytosolic GAPC (Moreno-García et al., 2022). Overall, this observation points towards an increase in the NADH/NAD^+^ ratio, due to re-routing of malate from the mitochondria to the cytosol.

### Light intensity modulates transcriptomic reprogramming

To capture shifts in transcriptional reprogramming at the dark-light transition, we performed RNA-seq on Col-0 and *mdh1xme1xme2.1* grown under SD-LL or SD-NL. Genotype-dependent expression differences were modulated profoundly by irradiance. Principal component analysis (PCA) showed that under LL, wild type and the triple mutant diverged most strongly one hour after onset of illumination (‘lights-on’; +1), whereas under NL a comparable separation was evident one hour before ‘lights-on’ (-1) (Fig. 6A). Consistent with this phase shift, the triple mutants showed the strongest transcriptional response to light onset under LL, with 160 low-abundance and 258 higher-abundance transcripts (Fig. 6B). By contrast, under NL the largest genotype-dependent differences occurred at the end of the night, comprising 611 down- and 959 up-regulated transcripts (Fig. 6B). Together, these data indicate that irradiance rephases the mutant transcriptional phenotype: LL reveals a dawn-associated constraint consistent with impaired establishment of photosynthetic/redox homeostasis that coincides with chlorosis and slow growth, whereas NL largely compensates daytime function and shifts the major molecular signature to the end of the night, suggesting altered nocturnal metabolism and/or anticipatory dawn programming. This supports a conditional, environment-dependent metabolic limitation rather than a constitutive developmental defect.

**Figure 6.**
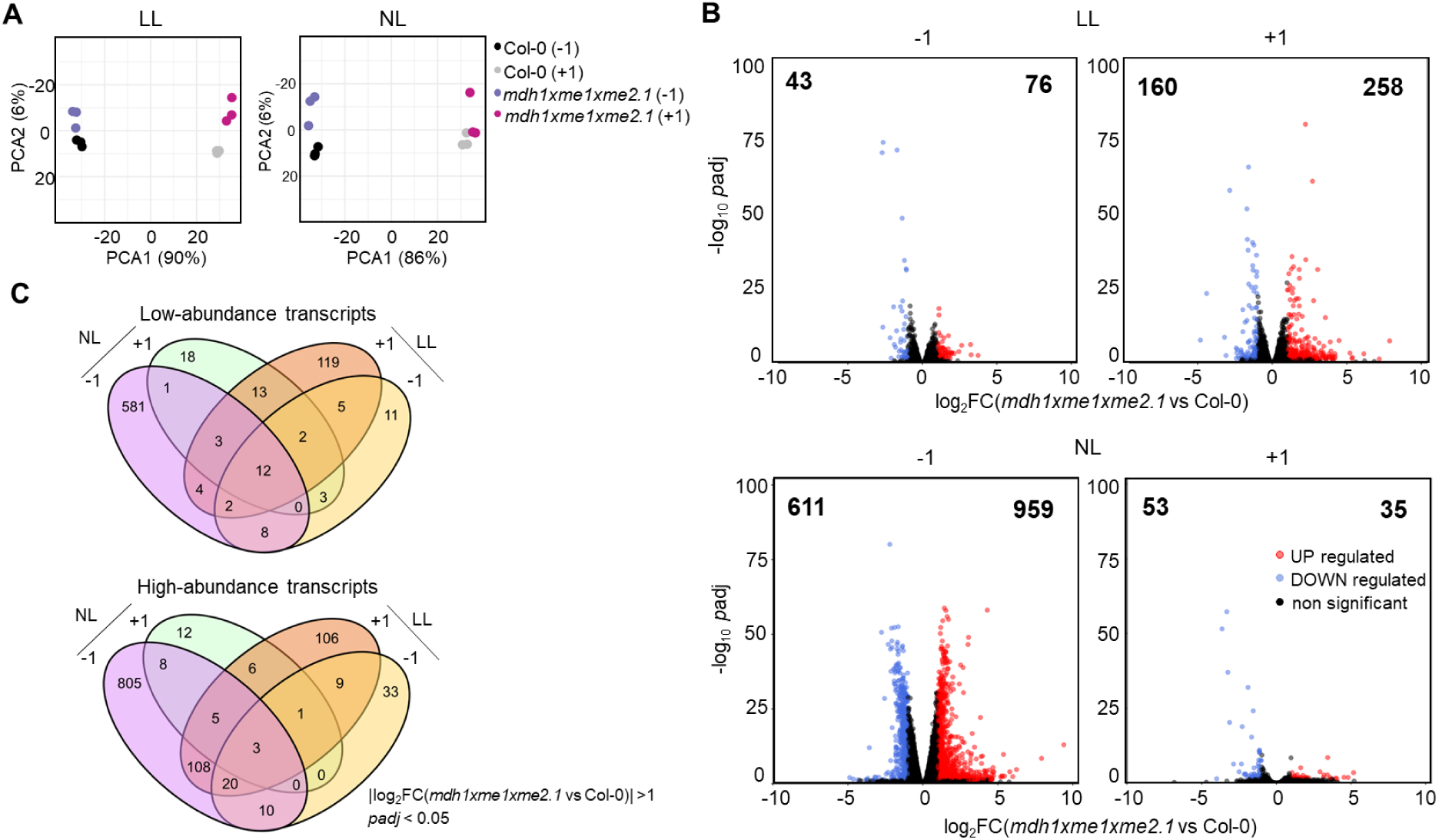
Differential gene expression during the dark-light transition. **A)** Principal component analysis of RNAseq data from whole rosettes grown at low light (LL) and (normal light (NL). For both light intensities, samples were collected 1 h before lights-on (-1) and 1 h after lights-on (+1). **B)** Volcano plots show genes up- and down-regulated in *mdh1xme1xme2.1* relative to wild type. Genes are considered differentially expressed at |log_2_FC |> 1 and a *p*adj-value < 0.05 (Suppl. Data 1). Differential expression was assessed with DESeq2 package in R studio (n = 3 per genotype and condition). Numbers in the upper left and right corners indicate the counts of up- and down-regulated transcripts, respectively. **C)** Venn diagram showing common and condition-unique transcripts in *mdh1xme1xme2.1* at LL and NL and -1 and +1 (high abundance transcripts: log2FC > 1, padj < 0.05; low abundance transcripts: log2FC < −1, padj < 0.05; n = 3 per genotype and condition).

To focus on condition-specific transcriptional responses, we identified transcripts uniquely up-or down-regulated in each growth regime (Fig. 6C). The resulting condition-specific gene sets were then subjected to Gene Ontology (GO) enrichment analysis using STRING v12.0 (Szklarczyk et al., 2023) to pinpoint the most strongly affected biological processes (Suppl. Fig. 5 and Suppl. Data 1).

Under LL, one hour after lights-on (+1), *mdh1xme1xme2.1* displayed 119 transcripts that were uniquely repressed under this condition. GO enrichment analysis indicated that these genes are predominantly associated with microtubule-based processes, cell cycle processes, cell division, and cytoskeleton organization (Supp. Fig. 5; Suppl. Data 1). This set includes multiple kinesin-like proteins as well as microtubule-associated proteins that are required for plant growth and mitotic progression (Li et al., 2012; Krtková et al., 2016). The coordinated repression of cell-cycle and division-related programs is consistent with the delayed development and reduced biomass accumulation of *mdh1xme1xme2* under LL. Among the lowest abundance genes are those associated with the cellular response to iron starvation, including the basic helix-loop-helix transcription factors *bHLH100* (At2g41240; 28-times), *bHLH38* (At3g56970; 21-times), and *bHLH39* (At3g56980; 9-times). These members of the bHLH Ib subgroup activate iron-deficiency responses and iron uptake in *A. thaliana* (Yuan et al., 2008; Wang et al., 2013). Iron is critical for photosynthetic performance because it serves as an essential cofactor in both photosystems (Ben-Shem et al., 2003; Umena et al., 2011). Accordingly, iron limitation reduces photosynthetic capacity and can alter chloroplast development, including chloroplast morphology and the abundance of photosynthetic complexes (Spiller and Terry, 1980; Larbi et al., 2004). The low expression of iron-starvation response genes is therefore consistent with the reduced photosynthetic efficiency and altered chloroplast morphology observed in *mdh1xme1xme2* under LL. Transcripts associated with carbohydrate biosynthetic process were also lower abundant in *mdh1xme1xme2.1* under LL. Many of these genes encode enzymes involved in cell-wall formation, including cellulose synthases and xyloglucan endotransglucosylase/hydrolase proteins that build and remodel key components of the expanding cell wall in growing tissues. The coordinated low expression of this gene set is consistent with reduced investment in cell-wall biosynthesis and provides a plausible molecular correlate of the growth inhibition observed specifically under LL. Notably, this GO category also includes *RBCS1A* (ribulose bisphosphate carboxylase small subunit 1A; At1g67090) which was expressed 3-times lower than in the wild type. *RBCS1A* is the most highly expressed member of the *RBCS* family in wild-type plants, and loss-of-function mutants show reduced Rubisco abundance and impaired photosynthesis, (Izumi et al., 2012), consistent with the diminished photosynthetic performance of *mdh1xme1xme2.1* under LL.

The triple mutant also exhibited 106 transcripts that were uniquely induced in SD-LL one hour after lights-on (+1). GO enrichment identifies response to phosphate starvation as the dominant category (Suppl. Fig. 5), indicating activation of a Pi-starvation-like program. In line with this, *SPX3* (At2g45130), SPX domain-containing protein 3, a positive regulator of Pi-starvation acclimation, was about 14-times higher expressed, and *PS2* (At1g73010), encoding a cytosolic pyrophosphatase that can mobilize Pi from PPi (May et al., 2011), increases about 7-times. The induction of two monogalactosyldiacylglycerol synthase, *MGD2* (At5g20410; (∼3.5-times) and *MGD3* (At2g11810; ∼5-times) further supports engagement of the canonical lipid- remodeling response associated with Pi limitation (Awai et al., 2001; Kobayashi et al., 2004; Kobayashi et al., 2009). Importantly, Pi scarcity is tightly coupled to photosynthetic performance and internal Pi redistribution (Schachtman et al., 1998; Kobayashi et al., 2009). Thus, even if external Pi is not limited, the observed module is consistent with a perceived Pi imbalance that could plausibly contribute to the SD-LL phenotype by constraining chloroplast metabolism at the onset of the light period. One mechanistic link is carbon/organic-acid metabolism: Pi acquisition in soil frequently involves organic-acid exudation (e.g., malate/citrate) to mobilize sparingly available phosphate (Ryan et al., 2001). This raises the possibility that compromised mitochondrial malate utilization reduces the availability or flexibility of organic-acid pools, thereby limiting Pi-foraging capacity and/or the ability to buffer cytosolic Pi/PPi homeostasis under the energy-constrained regime. Beyond the Pi-starvation module, the remaining uniquely induced processes clustered into responses to diverse stressors (Suppl. Fig. 5), suggesting a broader stress-acclimation state triggered specifically in SD-LL at dawn.

Under LL, one hour before lights-on (-1) only 11 transcripts were uniquely lower abundant in the triple mutant, whereas 33 were uniquely higher abundant (Fig. 6; Suppl. Data 1). Most changes were modest (approximately 2-3-times). The main outlier was At3g57520 (∼6-times higher), encoding an alkaline α-galactosidase implicated in raffinose hydrolysis to sucrose (supporting carbon export) and thylakoid galactolipid turnover during senescence (Gaudreault and Webb, 1986; Bachmann et al., 1994; Carmi et al., 2003; Lee et al., 2009) . Notably, the iron-homeostasis signature seen after lights-on was already evident pre-dawn: *ferritin 1*,*FER1* (At5g01600), was ∼2-times higher, whereas *bHLH101* (At5g04150), a bHLH Ib factor promoting iron uptake, was ∼3.5-times lower in the triple mutant (Pu and Liang, 2023).

Under SD-NL, *mdh1xme1xme2.1* showed its strongest transcriptome remodeling before dawn (-1), with hundreds of transcripts uniquely higher and lower abundant (Fig. 6; Suppl. Data 1). The dominant themes point to broad reprogramming of protein production and turnover, accompanied by shifts in growth-related hormone signaling and changes in chromatin/transcriptional regulation, consistent with extensive regulatory adjustment during the night. In parallel, transcripts linked to mitochondrial import and respiration (and associated stress buffering) are elevated, while photosynthesis-related expression remains compatible with an essentially normal phenotype. Around dawn, the mutant also showed a targeted repression of sulfur acquisition and sulfur-intensive defense/detox pathways, alongside induction of a small set of stress-associated genes, suggesting resource reallocation at day onset. Overall, NL condition appears to allow compensatory transcriptional acclimation that buffers mitochondrial malate disruption, in contrast to the failure of compensation under lower-energy light regime.

### Light intensity-dependent proteome shifts reveal genotype-specific responses

Given the pronounced physiological defects caused by combined disruption of MDH1 and ME under low-energy conditions, and to complement the transcriptomic signatures observed across light regimes, we performed a comprehensive proteomic analysis on leaves harvested 1.5 h after lights-on from Col-0 and *mdh1xme1xme2.1* and *mdh1xme1xme2.2* grown under SD at LL and NL.

PCA showed a strong separation between wild type and the triple mutants under LL, whereas this separation was reduced under NL (Suppl. Fig. 6A), consistent with a partial proteomic recovery under non-limiting light condition. iBAQ-based protein quantification (Schwanhäusser et al., 2011), grouped by subcellular localization using SUBAcon (Hooper et al., 2014), revealed a LL-specific redistribution of protein allocation: both mutants have a reduced relative abundance of plastid-targeted proteins, matching the chloroplast structural defects, and a relative increase in cytosolic, nuclear, peroxisomal, and mitochondrial proteins (Suppl. Fig. 6B). Under NL, this compartmental shift was largely absent, aside from a reduction in chloroplast proteins in *mdh1xme1xme2.2*.

To compare plastid proteomes without bias from relative reduced plastid protein content in the mutants, we applied separated normalization for plastid and non-plastid proteins. This enabled condition-specific identification of altered processes in *mdh1xme1xme2* under SD at both LL and NL within each compartment. Protein groups were then assigned MapMan categories (Thimm et al., 2004), and pathway impact was summarized by counting significantly changed protein groups per process. The most pronounced proteomic differences between *mdh1xme1xme2* and Col-0 were observed under LL (Supp. Fig. 7). In LL-grown *mdh1xme1xme2.1*, 177 plastid and 179 non-plastid protein groups were less abundant than in Col-0, whereas 232 plastid and 175 non-plastid groups were more abundant (Supp. Fig. 7A). In contrast, under NL the number of differentially abundant proteins was substantially lower (Suppl. Fig. 7B), consistent with a partial normalization of the proteome at higher irradiance. *mdh1xme1xme2.2* showed a similar pattern (Supp. Fig. 7).

### Chloroplast proteome remodeling in mdh1xme1xme2 under LL favors stress acclimation over photosynthetic capacity

Under LL, the chloroplast proteome of *mdh1xme1xme2* was extensively remodeled relative to Col-0 (Suppl. Fig. 8; Suppl. Data 2), consistent with the strong phenotype under this regime. A prominent signature is a rebalancing of plastid protein homeostasis: protein-metabolism categories account for a large fraction of the differentially abundant plastid protein groups, with a clear depletion of plastid translation capacity. Specifically, most of the reduced protein-metabolism groups correspond to plastid ribosomal subunits, whereas proteins associated with proteolysis and import/translocation accumulate. This pattern suggests that LL imposes a state where the mutant chloroplast prioritizes turnover/quality control over net biosynthesis, likely limiting the ability to build and maintain the photosynthetic apparatus.

Consistent with this, plastid proteins associated with photosynthesis-related were extensively remodeled (Suppl. Fig. 8; Suppl. Data 2). A prominent change was the reduction in Rubisco abundance; multiple small-subunit isoforms (RBCS1A, RBCS1B, RBCS2B) and the large subunit (RBCL) were each decreased by more than 2-times in the mutant. This decrease provides a direct and plausible molecular explanation for the lower CO**_2_** assimilation and slower growth under LL, since Rubisco content is a major determinant of carbon fixation capacity (Lobo et al., 2019). Beyond Rubisco, several core components of Photosystem I (PSI) and light harvesting complex I (LHCI) were reduced, accompanied by a broad decrease in NADH dehydrogenase-like (NDH) complex subunits. Because NDH-dependent cyclic electron flow supports ATP production and alleviates stromal over-reduction, functions that are particularly important under low light and after dark adaptation (Yamori et al., 2011; Kou et al., 2015; Martín et al., 2015; Basso et al., 2020), the concomitant loss of NDH and PSI depletion suggests a diminished capacity in the mutant to adjust the ATP/NADPH balance and to buffer redox pressure under LL. Several Photosystem II (PSII) core/auxiliary proteins were also less abundant (e.g., PSBQ-2, PSBR, PSBO-2), and the thylakoid phosphatase TAP38, which is required for efficient light harvesting complex II (LHCII) dephosphorylation and redistribution of excitation-energy between PSII and PSI (Pribil et al., 2010; Shapiguzov et al., 2010), was strongly reduced. Collectively, these changes point to chloroplasts that are both limited in assembling photosynthetic complexes and impaired in optimizing excitation balance, features likely to intensify chlorosis and depress photosynthetic performance under energy-limited conditions.

Notably, the proteins that increased within the photosynthesis category were biased toward photoprotection and PSII maintenance. About half were PSII-associated, including LHCII components (e.g., LHCB isoforms linked to grana organization (Albanese et al., 2020)), PsbS/NPQ4 (consistent with elevated NPQ; (Li et al., 2000; Peterson, 2005; Johnson and Ruban, 2011)), and PSII repair/assembly factors (e.g., PsbP-like proteins and MPH2; (Ishihara et al., 2007; Liu and Last, 2017)). This pattern suggests that under LL the mutant shifts resources toward excitation dissipation and PSII maintenance rather than photochemical output, enhancing protection at the expense of productivity. Chloroplast metabolism also showed compensatory shifts consistent with reduced Rubisco: several Calvin-cycle proteins increased, including plastid Glyceraldehyde-3-phosphate dehydrogenase (GAPDH) subunits (GAPA1, GAPA2, GAPB), together with elevated Rubisco assembly factors (e.g., RAF1 isoforms), suggesting an attempt to sustain carbon flux and rebuild/stabilize the Rubisco holoenzyme (Hauser et al., 2015; Whitney et al., 2015).

At the same time, plastid amino-acid and nitrogen-assimilation proteins were rebalanced, with reduced abundance of NADH-dependent glutamine-2-oxoglutarate aminotransferase (GOGAT, At5g53460) and glutamine synthetase 2 (GLN2, At5g35630), pointing to altered ammonium assimilation routes and a broader disturbance in C-N coordination under LL.

Finally, the most prominent stress-related plastid signature point to altered iron handling and enhanced ROS detoxification. Ferritins accumulated, most notably FER1, with FER3/FER4 also increased, together with multiple H_2_O_2_-scavenging and redox-homeostasis proteins, including stromal ascorbate peroxidase (APX) and related factors. Because ferritins buffer free iron and thereby limit iron-catalyzed ROS formation (Briat and Lobréaux, 1997; Murgia et al., 2007; Ravet et al., 2009), this coordinated response is consistent with elevated oxidative pressure in LL-grown mutant chloroplasts, plausibly stemming from impaired electron transport and redox balancing.

Overall, the LL plastid proteome indicates a shift away from chloroplast translation and photosynthetic capacity toward proteome maintenance, photoprotection/repair, and iron/ROS management. This provides a coherent molecular rationale for why LL uniquely reveals chlorosis and growth defects: the mutant appears to adopt a protective, stress-acclimation state that preserves chloroplast integrity but constrains carbon gain and biomass accumulation under energy-limited conditions.

### Cytosolic proteome changes in mdh1xme1xme2 indicate nitrogen remobilization and stress acclimation under LL

Under LL, the cytosolic proteome of *mdh1xme1xme2* was strongly remodeled (Supp. Fig. 9; Suppl. Data 2), with most differentially abundant protein groups assigned to protein metabolism, carbohydrate metabolism, and amino-acid metabolism. Within protein metabolism, decreased proteins were enriched for biosynthetic functions, whereas increased proteins were biased toward protein modification. Carbohydrate metabolism was also affected: reduced proteins included enzymes linked to carbohydrate breakdown and nucleotide-sugar biosynthesis. Because nucleotide sugars supply activated donors for cell-wall polysaccharides, glycoproteins, and glycolipids (Reiter and Vanzin, 2001; Seifert, 2018), their decrease is consistent with reduced anabolic investment and slower growth under LL.

A key feature of the amino-acid category was a switch from nitrate assimilation toward nitrogen recycling. Glutamine-dependent asparagine synthase 1 (ASN1) accumulated strongly (∼7-times), consistent with its role as a C/N-responsive, dark-inducible enzyme promoting N remobilization when carbon is limiting and during senescence-like states (Lam et al., 1994; Nozawa et al., 1999; Fujiki et al., 2001). Pyruvate Phosphate Dikinase (PPDK) also increased (∼4-times), supporting broader rewiring of carbon-nitrogen flux typical of remobilization programs (Lin and Wu, 2004). In contrast, nitrate reductase 2 (NIA2) decreased (∼2-times), consistent with reduced nitrate assimilation capacity; NIA2 contributes most to nitrate reductase activity in Arabidopsis shoots (Wilkinson and Crawford, 1993). In parallel, stress/defense proteins were enriched, including mannose-binding lectins and phospholipase 2A (PLA2A), alongside strong accumulation of oxidative/xenobiotic stress markers such as glutathione transferases, GSTF6 and GSTF7 (∼6-times) (Sappl et al., 2009; Kouno and Ezaki, 2013).

Overall, the LL cytosolic proteome supports a shift of *mdh1xme1xme2* toward a stress-acclimated, remobilization-oriented state, marked by reduced nitrate assimilation and nucleotide-sugar-dependent biosynthesis, and increased detoxification and defense capacity.

### Mitochondrial proteome remodeling in mdh1xme1xme2 under LL points to increased amino-acid catabolism and altered transport capacity

Under low light (LL), the mitochondrial proteome of *mdh1xme1xme2* shifted toward sustaining respiration under limited photosynthetic carbon supply and impaired malate flux. Several TCA cycle-associated proteins increased in abundance, although only IDH (isocitrate/isopropylmalate dehydrogenase; At5g14590) showed a significant rise (Fig. 7, Suppl. Data 2). Notably, about one third of the upregulated mitochondrial proteins were linked to amino acid metabolism, predominantly catabolic pathways. This pattern suggests a compensatory strategy in which amino acid degradation supplies carbon skeletons and reducing equivalents to maintain respiratory activity when photosynthetic input is restricted (Fig. 7, Suppl. Data 2). Within this module, enzymes associated with stress-linked nitrogen and carbon reallocation accumulate. Arginase isoforms (ARGAH1 and ARGAH2) increased ∼2-times, consistent with activation of arginine turnover during stress (Brauc et al., 2012). Glutamate dehydrogenases (GDH1 and GDH2) were also elevated (∼3-times), supporting enhanced interconversion between glutamate and 2-oxoglutarate, a reaction frequently engaged under carbon limitation to feed the TCA cycle while releasing ammonia (Miyashita and Good, 2008; Fontaine et al., 2012). In addition, the increased abundance of Methylcrotonoyl-CoA carboxylase subunit alpha (MCCA; ∼4-times) and isovaleryl-CoA dehydrogenase (IVDH; ∼2-times) suggests use of branched-chain amino-acid (BCAA) to supply carbon and electrons, a response commonly observed under carbon scarcity and senescence-like metabolic states (Che et al., 2002; Araújo et al., 2010; Engqvist et al., 2011; Ding et al., 2012). Together, these changes indicate that LL-grown mutants partially compensate for impaired malate metabolism by mobilizing amino acids as alternative respiratory substrates.

**Figure 7.**
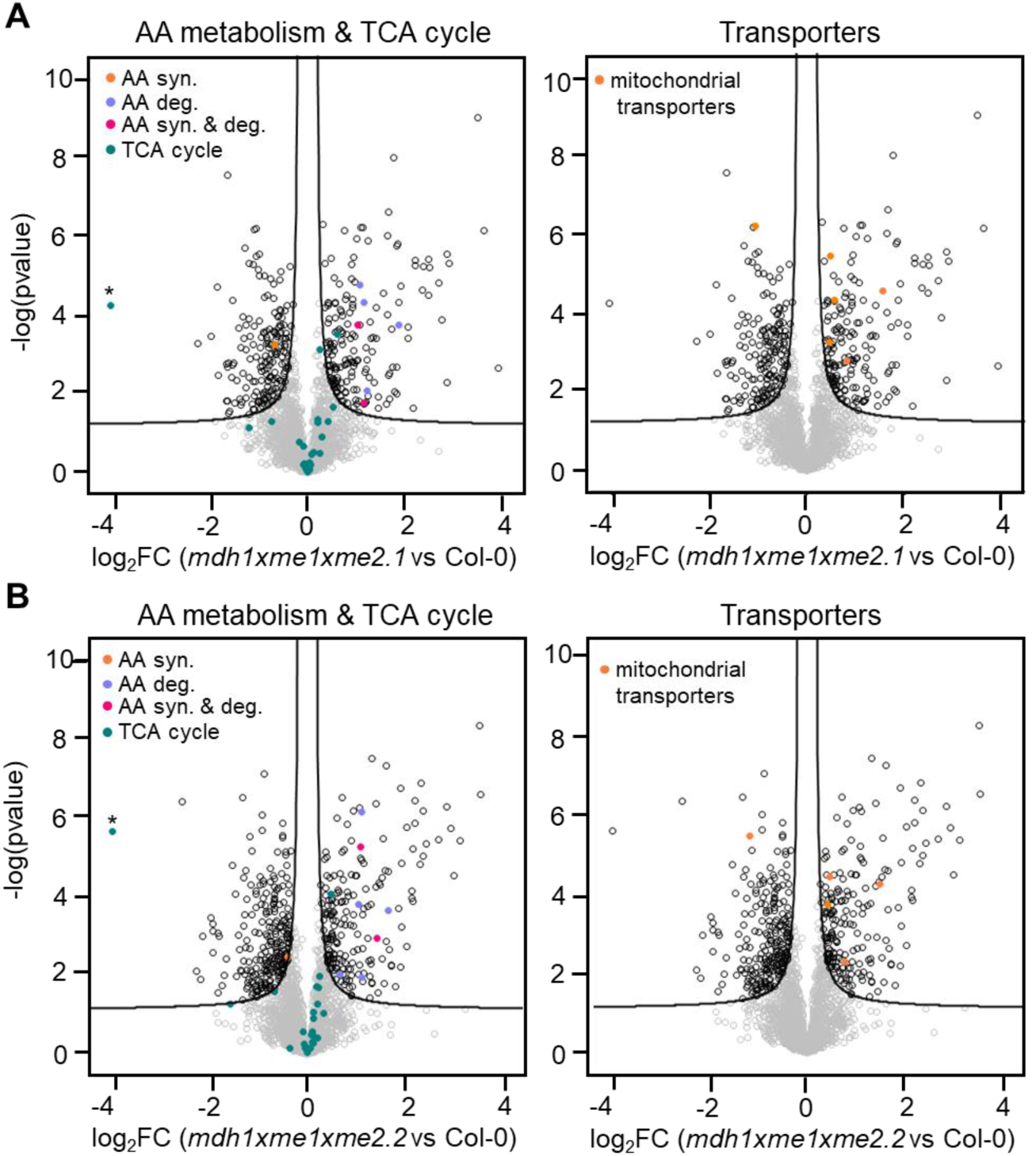
Differential abundance of mitochondrial protein groups in the *mdh1xme1xme2* under short days and low light. After normalization, log2-transformed LFQ values of protein groups not assigned to plastid by the SUBAcon algorithm were used to assess differences in the mitochondria proteome between **(A)** *mdh1xme1xme2.1* and Col-0 and **(B)** *mdh1xme1xme2.2* and Col-0 by statistical testing employing five independent biological replicates. Protein groups were classified by MapMan annotation in levels based on their biological function. The complete list of categories is provided in Supplemental Data 2. Statistical thresholds were set at FDR ≤ 0.05 and S₀ = 0.1. Protein groups located in the areas above the black lines fulfil the significance threshold limits (FDR ≤ 0.05; S_0_ = 0.1). syn: synthesis, deg: degradation, AA: amino acid, TCA: tricarboxylic acid. Protein group identified as MDH is marked by an asterisk, however as previously mentioned, no unique peptides of mitochondrial MDH1 were detected in *mdh1xme1xme2* mutants. Given the abundance of this enzyme within the mitochondrial compartment, which yields unique peptides in the WT samples and considering the changes in MDH1 transcript levels, the absence of unique peptides in the mutant lines is thus a strong indicator for the absence of this protein in these lines. LL: low light.

Transport functions also emerged as a prominent mitochondrial signature under LL (Fig. 7, Suppl. Data 2). Proteins associated with mitochondrial import were increased, including metaxin (∼3-times) and TOM20-2 (∼2-times), indicating enhanced protein import capacity and proteome remodeling. In contrast, ADNT1 (adenine nucleotide transporter 1), which mediates ATP/AMP exchange across the inner mitochondrial membrane, decreased by ∼2-times (Palmieri et al., 2008).

Overall, the LL mitochondrial proteome reflects a shift toward amino acid-supported respiration and restructured transport capacity, reinforcing the view that under energy-limited conditions the triple mutant mobilizes alternative substrates and remodels the organelle to sustain mitochondrial function.

### Cell wall biosynthesis and turnover are rebalanced in mdh1xme1xme2 grown under SD and LL

Under LL, cell-wall metabolism emerges as a major non-plastid target in *mdh1xme1xme2* (Supp. Fig. 10; Suppl. Data 2). Proteins linked to xyloglucan and cellulose deposition were reduced. Xyloglucan endotransglucosylase/hydrolases (XTH7 and XTH32) decreased ∼3-times, while the primary-wall cellulose synthases CESA1 and CESA3 were also lower in abundance suggesting reduced cellulose synthase complex activity and constrained cell elongation (Arioli et al., 1998; Persson et al., 2007). Conversely, several glycoside hydrolases, including β-galactosidases and β-xylosidases, accumulated, consistent with increased carbohydrate remobilization and cell wall turnover under sugar-limited or senescence-like conditions (Lee et al., 2009).

Taken together, the LL proteome indicates that low energy availability shifts *mdh1xme1xme2* away from cell wall construction toward enhanced degradation and remodeling. This reallocation likely contributes to the reduced growth and biomass accumulation observed under LL, consistent with a broader transition from growth to maintenance and resource recycling.

### Low light triggers C/N imbalance and ammonium accumulation in *mdh1xme1xme2*

The proteomic profile, combined with impaired photosynthetic performance under LL, indicates altered carbon-nitrogen (C/N) homeostasis in *mdh1xme1xme2*. To assess this, total carbon and nitrogen contents were measured in plants grown under SD at LL or NL.

Under LL, the triple mutant exhibited lower leaf C/N ratio than wild type, with the largest differences at the end of the night and at the end of the day (Fig. 8A-C). This shift reflects both reduced total carbon and increased total nitrogen, most prominently towards the end of the light period, indicating a progressive imbalance over the diel cycle (Fig. 8A-C). In contrast, no genotype-dependent changes in C/N were detected under NL (Suppl. Fig. 11), suggesting that the C/N phenotype is conditional on carbon limitation imposed by LL. Consistent with an altered nitrogen status under LL, NH_4_^+^ accumulated more strongly after light onset (+1) in the triple mutant lines compared with wild type, whereas no such increase was observed under NL (Suppl. Fig. 11).

**Figure 8.**
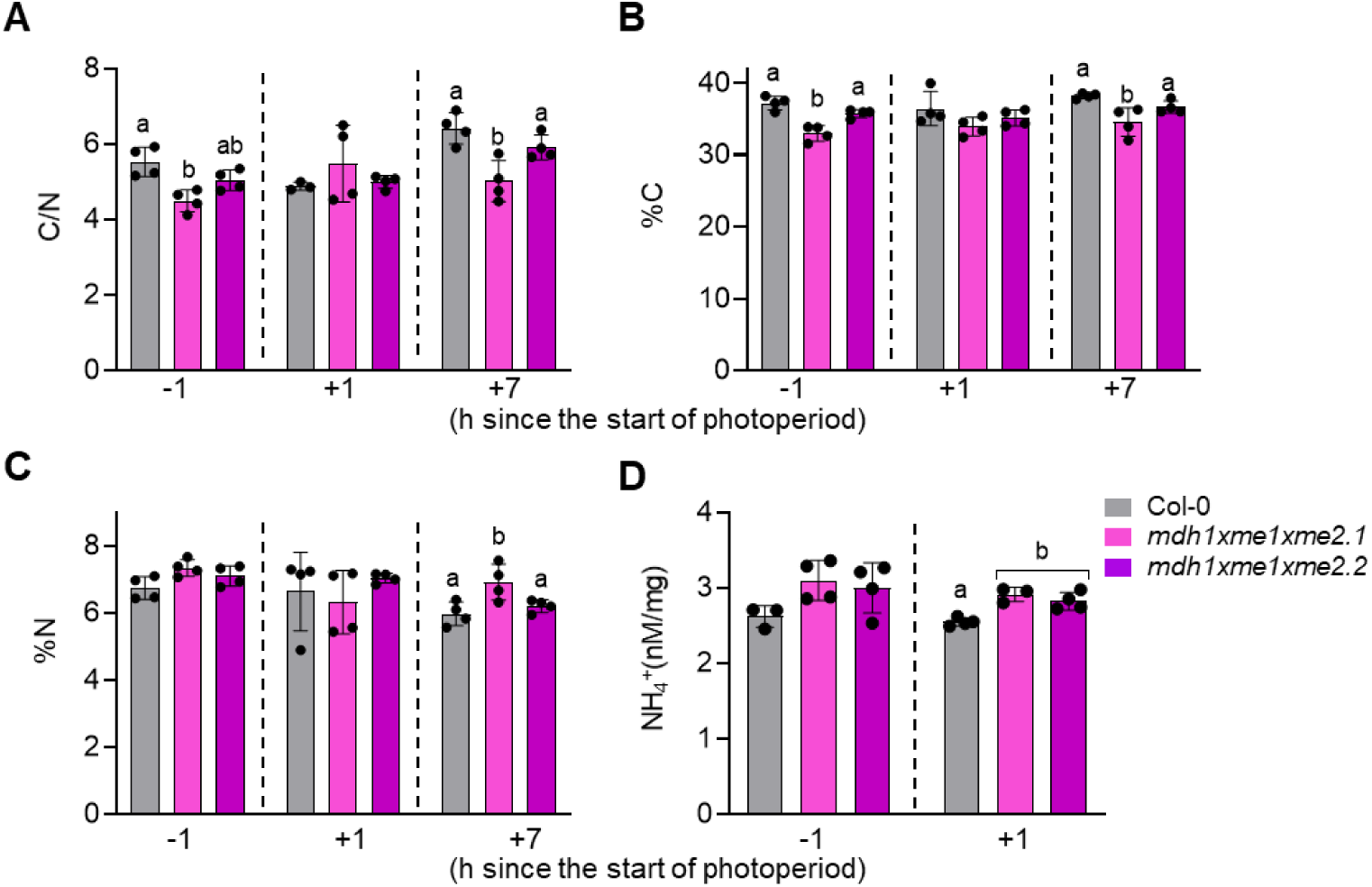
C/N ratio, total C, total N and ammonia in plants grown in SD and LL. **A)** Carbon-to-nitrogen ratio (C/N ratio), calculated as total carbon divided by total nitrogen. **B)** C content (%C) and **C)** N content (%N) expressed as percentage of dry weight in samples harvested 1 h before light onset (-1), one hour after light onset (+1), and one hour before light offset (+7). **D)** Leaf total ammonia (NH_4_^+^) concentrations measured one hour before light onset (-1) and one hour after light onset (+1). Data represent means ± SD of 3-4 biological replicates per line. Different letters indicate significant differences (*p* < 0.05) according to ANOVA followed by Tukey’s post-hoc test. LL: low light; SD: short day.

Proteomics further supports a shift in ammonium-associated metabolism: both GDH subunits increase in abundance in the triple mutants under LL (Suppl. Data 2). Although GDH can operate bidirectionally, *in vivo* mitochondrial GDH is commonly associated with glutamate deamination, providing 2-oxoglutarate to sustain the TCA cycle under carbon scarcity or extended darkness, while concomitantly generating NH_4_⁺ (Fontaine et al., 2012). Thus, the induction of GDH in *mdh1xme1xme2* under LL and long nights may reflect an increased reliance on the GDH shunt to support respiratory carbon flux, with the trade-off of elevated NH_4_⁺ release.

Functionally, a combination of lower carbon availability, relative nitrogen excess, and enhanced NH_4_⁺ accumulation after dawn is consistent with an NH_4_⁺ stress scenario that could contribute to the growth defect under LL. Excess NH_4_⁺ in photosynthetic tissues can impose a substantial energetic burden for reassimilation and promote cellular acidification, ultimately restricting growth through acidic stress and energy depletion (Hachiya et al., 2021). Moreover, increased chloroplast NH_4_⁺ has been linked to photosynthetic inhibition and PSII photodamage (Drath et al., 2008), providing a plausible mechanistic connection between NH_4_⁺ accumulation and the impaired photosynthetic performance observed in the triple mutants under LL. Taken together, our data indicate that *mdh1xme1xme2* fails to maintain C/N homeostasis under carbon-limiting conditions, leading to NH_4_⁺ accumulation and activation of GDH-associated metabolism, which likely exacerbates stress on growth and photosynthetic function.

### Malate metabolism defect in *mdh1xme1xme2* triggers energy-dependent carbon and amino acid reprogramming

To assess the metabolic consequences of impaired mitochondrial malate metabolism in the triple mutants, we performed GCMS-based metabolite profiling of rosettes from plants grown (i) in SD at LL, and (ii) under SD and LD at NL. A total of 35 metabolites were measured.

Across all conditions tested, malate abundance was higher in *mdh1xme1xme2* than in Col-0 at both the end of the night and the end of the day (Fig. 9). Persistent malate accumulation mirrors a sustained disruption of malate turnover, independent of photoperiod or light intensity, consistent with the loss of mitochondrial malate-converting activities. However, the physiological and downstream metabolic consequences of this defect depended on the energy regime. Under SD-LL, pyruvate levels were reduced at the night-to-day transition but return to wild-type levels by the end of the day (Fig. 9). This profile suggests increased pyruvate consumption during the long night, likely to support mitochondrial metabolism when carbon availability is limiting. Under SD-NL, pyruvate recovered to wild-type levels within 1 h after light onset (+1), indicating that higher irradiance enables more rapid replenishment of pyruvate, likely via improved daytime carbon assimilation. Consistently, under LD-NL, total pyruvate levels in the mutants were indistinguishable from wild type (Fig. 9). Alanine, a potential alternative source of pyruvate, displays a strong diel signature. Across all conditions, alanine accumulated to about double the levels in the mutants at -1 (Fig. 9). Under SD-LL, alanine dropped at +1 and reverted to wild-type levels, whereas under SD-NL and LD-NL it remained elevated relative to wild type after illumination (Fig. 9). This light-dependent decline specifically under LL is consistent with preferential alanine utilization at dawn, when carbon and energy supply are most limiting.

**Figure 9.**
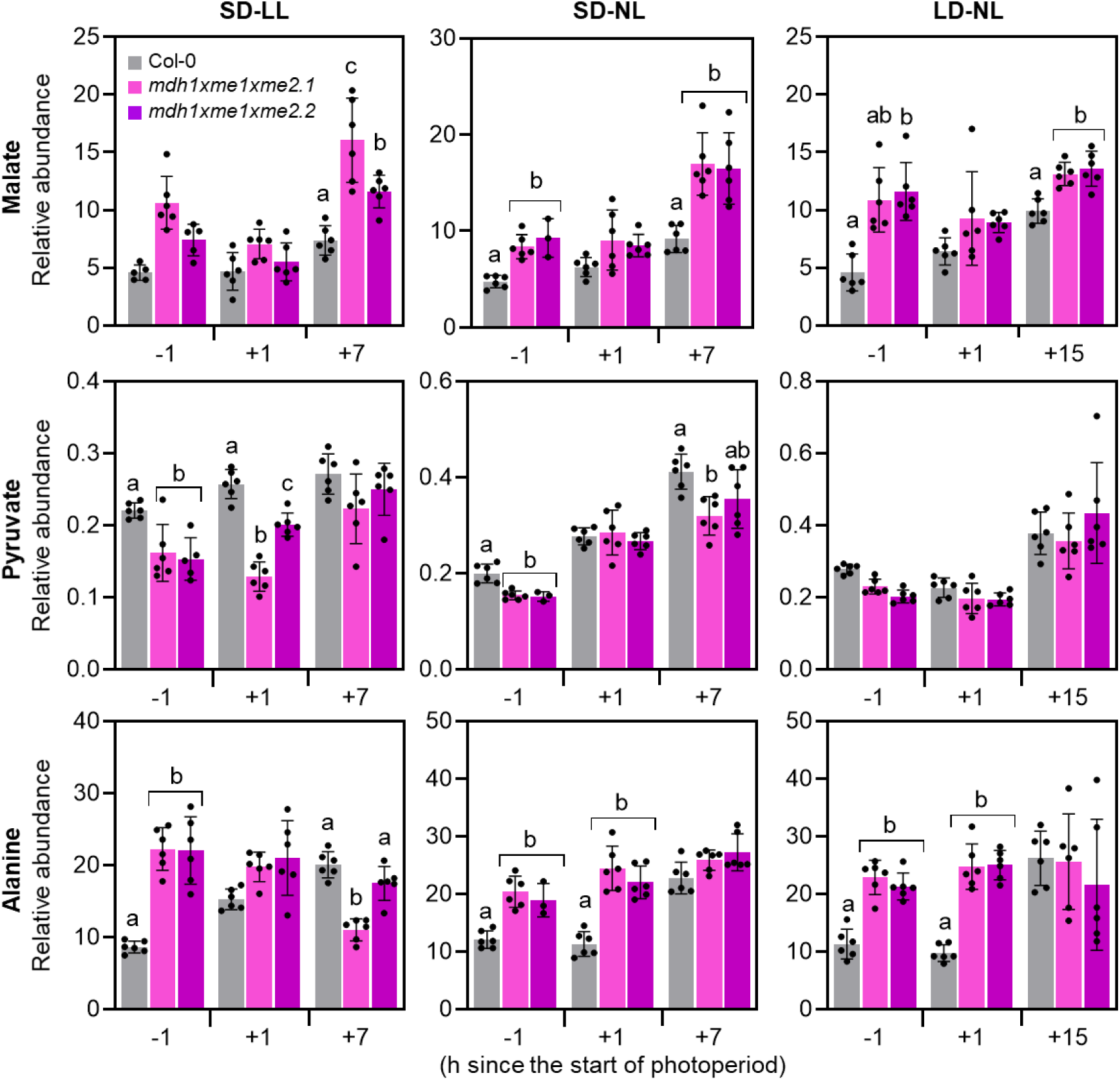
Relative concentrations of malate, pyruvate and alanine. Relative levels were quantified by GC-MS in leaves of the indicated genotypes grown under LL-SD, NL SD and NL LD. Samples were harvested 1 h before lights-on (-1 h), 1 h after lights-on (+1 h) and 1 h before lights-off (+7/+15 h). Bars show means ± SD (n = 6 independent biological replicates). Statistics were performed separately for each time point using one-way ANOVA followed by Tukey’s post-hoc test; bars that do not share a letter differ significantly at p < 0.05. LD: long day; LL: low light; NL: normal light; SD: short day.

The branched-chain amino acids (BCAAs), leucine, isoleucine and valine, catabolism is an important function of mitochondrial matrix metabolism (Diebold et al., 2002; Ishizaki et al., 2005; Schuster and Binder, 2005). BCAAs accumulated in *mdh1xme1xme2* under LL, with the strongest increase at the end of the night (-1) (Suppl. Fig. 12). Similar BCAA elevations have been reported in wild-type plants during prolonged darkness and are thought to reflect metabolic adjustments that help sustain mitochondrial function under low-energy conditions (Engqvist et al., 2011; Peng et al., 2015; Li and Hoppe, 2023). Notably, alpha-ketoglutarate (αKG), a central TCA intermediate and major amino-group acceptor in transamination reactions, was strongly affected. In *mdh1xme1xme2* under LL, αKG levels at the end of the light period were ∼6-fold lower than in wild type (Suppl. Fig. 12). Importantly, in Col-0, αKG levels at -1 were ∼4-times higher under LL than under NL (Suppl. Fig. 12), underscoring the strong adjustment of this metabolite in wild type during low-energy nights. Overall, the rest metabolites measured did not show a particular trend in *mdh1xme1xme2* mutants (Suppl. Data 3).

## Summary and Conclusions

Using Arabidopsis *mdh1xme1xme2* triple mutants lacking the predominant mitochondrial MDH1 and both NAD-ME subunits (ME1/ME2), we show that mitochondrial malate turnover becomes a key metabolic constraint specifically under energy-limited regimes. Under SD combined with LL, the mutants exhibit dwarfism, chlorosis, and delayed development, accompanied by impaired photosynthetic performance (reduced PSII efficiency and CO_2_ assimilation, elevated NPQ) and altered chloroplast ultrastructure. In contrast, increasing irradiance or extending the photoperiod largely restores growth and photosynthetic competence, while elevated CO_2_ provides a partial rescue, indicating that limited photosynthetic carbon input amplifies the consequences of impaired mitochondrial malate utilization.

Multi-omics profiling links this conditional phenotype to a failure to sustain coordinated organelle function when carbon supply is limiting. Under LL, the mutant transcriptome reveals a pronounced dawn-associated bottleneck with reduced expression of growth-related programs and induction of stress-associated responses. This is paralleled by extensive proteome remodeling: the chloroplast shifts away from biosynthetic investment (including reduced abundance of photosynthetic components) toward protein quality control, photoprotection, and redox/iron-stress management, consistent with constrained productivity despite active energy dissipation.

At the whole-plant metabolic level, disrupted mitochondrial malate metabolism drives a reconfiguration of carbon-nitrogen balance under low energy. The mutants display lowered C/N ratios and post-dawn ammonium accumulation, alongside signatures of increased nitrogen remobilization and reduced nitrate assimilation. Metabolite and proteome patterns further support compensatory fueling of mitochondrial metabolism through amino-acid-derived carbon, including accumulation of pyruvate-related amino acids and branched-chain amino acids and increased abundance of mitochondrial BCAA catabolic enzymes, together with strong perturbation of 2-oxoglutarate homeostasis.

Taken together, our results identify mitochondrial malate conversion capacity as a control hub that integrates respiratory energy supply with photosynthetic function and stabilizes carbon allocation and nitrogen metabolism when plants experience prolonged nights and low irradiance.

## Material and Methods

### Plant material

The plant lines used in this study include *Arabidopsis thaliana* L. Heynh. ecotype Columbia (Col-0, wild type), along with homozygous T-DNA insertion lines Sail-374-A02 (*me1*), Sail-291-C05 (*me2.1*), and SALK_131720 (*me2.2*) (Tronconi et al., 2008), as well as GABI_097C10 (*mdh1*; (Kleinboelting et al., 2012)) and SALK_126994 (*mdh2*) (Tomaz et al., 2010) (Suppl. Table 1). The T-DNA insertion sites were verified by sequencing using specific primers (Suppl. Table 2; Suppl. Fig 1A). Double and triple homozygous mutants were generated by genetic crosses. The triple mutants were produced by crossing *me1* with *mdh1*x*me2.1* and with *mdh1*x*me2.2*. Homozygosity was confirmed by PCR using gene-specific and T-DNA left border primers (Suppl. Table 2; Suppl. Fig 1A).

### Plant growth

Plants were grown in soil (Floradur ® Substrate Multiplication Seed B, Rüttger Abstohs Gärtner-Einkauf GmbH&Co.KG, Siegburg, Germany) at 22/19 °C (day/night) under either long-day (LD: 16 h light/8 h dark) or short-day (SD: 8 h light/16 h dark) photoperiods, with light intensities of 60 µmol photons m^-2^ s^-1^ (LL) or 120 µmol photons m^-2^ s^-1^ (NL) and relative humidity of 70 %. Illumination was provided by a combination of Osram HO 80W/840 and HO 80W/830 bulbs. Seeds were stratified at 4 °C for 3 days prior to transfer to growth conditions. For all the experiments, whole rosettes were harvested in SD at LL light intensity at 40 days after stratification (DAS), at NL at 30 DAS, and in LD at 23 days after stratification. If this set up does not apply in a given method, it is specified in its own section.

Growth under high CO_2_ was conducted in a cabinet (Bronsonclimate, Holland) with a concentration of CO_2_ of 1100 ppm for a high CO_2_ condition and 400 ppm for normal conditions. Plants were grown under LL in short day photoperiod. Dry and fresh weight was determined in 47-days-old plants.

### Oxford nanopore technology (ONT) sequencing

High molecular weight DNA was extracted from young *A. thaliana* plants using a modified CTAB-based method (Siadjeu et al., 2020). The Short Read Eliminator kit (Pacific Biosciences) was used to deplete short DNA fragments from the samples. Samples were prepared for nanopore sequencing following the SQK-LSK114 protocol starting with 1 µg of DNA. Sequencing was performed on R10.4.1 flow cells on a MinION Mk1B. POD5 files were subjected to basecalling with dorado v1.0.2 (https://github.com/nanoporetech/dorado) using the high accuracy model dna_r10.4.1_e8.2_400bps_hac@v5.2.0. Reads were aligned to the TAIR10 reference genome sequence with minimap v2.27-r1193 (Li, 2018) using default parameters for nanopore reads. Samtools v1.19.2 (Li et al., 2009) was used to convert SAM into BAM files for visualization in Integrative Genomics Viewer v2.7.2 (Robinson et al., 2011). Bedtools (Quinlan and Hall, 2010) was utilized to infer coverage for each genomic position (Pucker and Brockington, 2018). Loreta v1 (Pucker et al., 2021) was deployed for the investigation of T-DNA insertions and structural rearrangements. The sequences of pAC161 (AJ537514) and pCSA110 (McElver et al., 2001) served as bait for the discovery of T-DNA sequences. A dedicated Python scripts was applied for the genome-wide coverage analysis (available viaGitHub: https://github.com/bpucker/GKseq2).

### Leaf surface area

For the calculation of the leaf surface area and maximum diameter a sample size of n = 10 was used from each genotype. The growth process was visually documented throughout the plant development and under NL and LL in SD conditions. Each plant picture was then analyzed using the software ImageJ build 1.50e. The leaf surface areas of all plant lines were determinate using Rosette Tracker image analysis (De Vylder et al., 2012) and were then compared with each other using a two-way ANOVA followed by Tukey’s post-hoc test. Statistical analyses were performed with the software RStudio using the package “multicomp” (version:1.4-29; (Hothorn et al., 2008)) which compared the leaf surface area of each plant line with each time point (DAS), and the same was applied for the maximum diameter.

### RNA extraction and cDNA synthesis

Total RNA was extracted from rosette leaves of plants grown in SD under LL and NL harvested 40 DAS and 30 DAS, respectively, using the Universal RNA Purification Kit (Roboklon GmbH, Germany) according to the manufacturer’s instructions, and eluted in 30 µl of RNase-free distilled water. The harvest points were one hour before the light was on (-1), one hour after the light was on (+1) and one hour before the light was off (+7). RNA concentration and purity were determined using a NanoDrop Nabi™ UV/Vis spectrophotometer (MicroDigital Co. Ltd, South Korea), and RNA integrity was verified by agarose gel electrophoresis. Samples were stored at -20 °C until further use.

To remove potential genomic DNA contamination, RNA samples were treated with the Ambion DNA-free™ DNA Removal Kit (Thermo Fisher Scientific, USA). Subsequently, 2 µg of RNA was reverse-transcribed into complementary DNA (cDNA) using RevertAid Reverse Transcriptase (Thermo Scientific, USA) and Oligo(dT) primers, following the manufacturer’s protocol.

### Reverse-transcription quantitative PCR

Reverse transcription quantitative PCR (RT-qPCR) was conducted using the KAPA SYBR FAST qPCR Kit (KAPA Biosystems Inc., Switzerland) to quantify the expression levels of *MDH1*, *MDH2*, *ME1*, and *ME2* in cDNA samples derived from the different plant lines. Reactions were performed on a Mastercycler® ep realplex system (Eppendorf, Germany), and cycle threshold (Ct) values were obtained using the accompanying realplex software. Gene expression was normalized to the *ACTIN2* (*ACT2I,* At3g18780) gene, which served as an internal reference.

### Pulse amplitude modulation fluorometry

PAM (Pulse Amplitude Modulation) measurements were performed using the IMAGING-PAM Maxi version (Walz, Germany) with a PAR (Photosynthetically Active Radiation) of 81 µmol m^-2^ s^-1^. Plants had been dark-adapted for 20 min prior to each measurement. For each plant line, four biological replicates were analyzed, and five measurement points were selected from each replicate. Average values of five points from each biological replicate were used to calculate each photosynthetic parameter. The maximum quantum yield efficiency of PSII (Fv/Fm = [Fm-F0]/Fm) was taken as a measure of PSII intactness, quantum Yield of PSII (ΦPSII) indicates the amount of solar energy converted to chemical energy to be used by the plant cell, the electron transport rate (ETR) shows the speed of the electrons in the electron transport chain and the non-photochemical quenching (NPQ).

### Gas exchange analysis

CO_2_ assimilation was measured using the Licor-6800 portable photosynthesis system (LI-COR Biosciences, USA) as outline in (Giese et al., 2023). Whole intact plants were used for determining CO_2_ assimilation and five biological replicates were analyzed per line and condition. The measurements were performed over a period of 30 min, starting one hour after the light was on. Light was set to NL and LL and gas exchange rate was evaluated using 700 µl flow with 5 % leak and 0.2 kPa overpressure. The reference CO_2_ was 400 µmol and the RH (Relative Humidity) was set to 80 %. The fan speed was set to 10,000 rpm and the heat exchanger temperature to 24 °C. Assimilation curves were done applying different CO_2_ concentrations (300, 400, 600, 800, 1000, 1200 and 1600 ppm). A blank reading was performed under the same conditions and subtracted from each measurement to avoid possible interferences. Assimilation values were normalized to plant area which was quantified using the ImageJ 1.45 in java 1.6.018 Rosette Tracker software (De Vylder et al., 2012)

### Pigment analysis

To quantify photosynthetic pigments, whole rosettes from plants growing in SD under NL and LL were harvested one hour before the onset of the light period. For pigment extraction, 20-30 mg of plant tissue was homogenized using TissueLyserII (QIAGEN, Germany) for 30 seconds two times at a frequency of 30 Hz. During the first homogenization, 500 µl of 100 % acetone was added to each sample. An additional 500 µl of 100 % acetone was added during the second homogenization, and samples were stored overnight at -20 °C. The following day, samples were centrifuged at maximum speed for 5 minutes, and the supernatant was filtered through a 0.2 µm membrane (GE Healthcare). The filtrate was transferred to new 1.5 ml Eppendorf tubes and analyzed by High Performance Liquid Chromatography (HPLC) as previously described (Farber et al., 1997). Pigment concentrations were normalized to fresh weight (FW).

### Transmission Electron Microscopy

The fourth and fifth leaves from plants grown under SD conditions at LL were excised one hour before the onset of the light period and immediately fixed in a solution containing 75 mM cacodylic acid sodium buffer (Carl Roth GmbH, Germany) pH 7 using HCl, 2.5 % (w/v) glutaraldehyde, 0.1 % (w/v) sucrose, and 2 mM MgSO₄ for 4-5 hours at 4 °C in a desiccator. Following fixation, leaves were washed three times for 15 minutes each in 100 mM cacodylate buffer. Post-fixation was performed in 2% (w/v) osmium tetroxide (OsO_4_) for 2.5 hours, followed by three 10-minute washes in distilled water. Leaves were then dehydrated through a graded acetone series (15 %-50 %), with three 10-minute incubations at each concentration. Subsequently, samples were incubated for 1 hour in 75 % (v/v) acetone containing 1 % (v/v) phosphotungstic acid and 1 % (w/v) uranyl acetate, followed by three 15-minute incubations in 80 % (v/v), 90 % (v/v), and 100 % (v/v) acetone. After dehydration, samples were infiltrated overnight with a 1:3 mixture of acetone and ERL-4206 resin (10 % (v/v) vinyl cyclohexene dioxide). This was followed by 4.5 hours in a 1:1 acetone/ERL mixture, 4 hours in a 3:1 acetone/ERL mixture, and finally overnight in pure ERL resin at 70 °C.

Once embedded, leaf sections were cut using an ultramicrotome and mounted on copper mesh grids. For contrast enhancement, grids were stained for 5 minutes in 1% uranyl acetate, followed by 5 minutes in a solution of lead nitrate [Pb(NO_3_)_2_] and sodium citrate [Na_3_C_6_H_5_O_7_·2H_2_O], and finally treated with 1 M NaOH, with water washes between each step. Prepared grids were examined using a Zeiss EM 10 transmission electron microscope (1978 model). Ultrathin sections were stained with Azur II 1 % (w/v), Methylene blue 1 % (w/v) dissolved in sodium tetraborate 1% (w/v) and observed using fluorescence stereomicroscope with 40x magnification lens (Leica MZFLIII).

### Respiration measurements

O_2_ consumption was measured in plants grown in SD at LL and in NL using the MitoXpress Xtra Oxygen Consumption Assay (Agilent, USA) as established recently (Wagner et al., 2019) with modifications. A total of 1 mL of assay buffer (10 mM MES, 10 mM MgCl_2_, 10 mM CaCl_2_, 5 mM KCl, pH 5.8) was used to reconstitute a vial containing dried MitoXpress® Xtra probe (Agilent Technologies, Santa Clara, CA, U.S.A) to create a stock solution. Prior to measurement, plants were kept in darkness and individual leaf discs from full expanded leaves were excised and weighed to reach 5 mg. The discs were immediately incubated with 10 µl of MitoXpress Xtra reagent and 140 µl of assay buffer for 1 hour under gentle shaking. The plate was monitored in a CLARIOstar® plate reader pre-equilibrated at 25 °C in TR-F mode, measuring probe signal in each well every 30 s. Instrument TR-F settings were: excitation filter – TR, dichroic mirror – LP-TR, emission filter – 645-20 nm, Delay times of 30 µs and 70 µs (two windows). After signal stabilization, 100 µL of mineral oil pre-warmed at 37 °C was added to each well to seal the samples from ambient air. Phosphorescence lifetime values (LT) for each sample and measurement point were calculated in Excel using the following formula accordingly to the manufacturers protocol: LT= (t2-t1)/ln(F1/F2) where t1 and t2 are the delay times (30 and 70 µs) and F1 and F2 are the corresponding intensity signals. The resulting LT values for each assayed well were plotted and the area under the curve was used to calculate the relative oxygen consumption. The inverse LT value was used in the final curve figure for a more intuitive visualization.

### Generation of Peredox-mCherry sensor lines and confocal imaging of cytosolic NAD redox dynamics

The Peredox-mCherry construct, which enables cytosol-specific expression of the biosensor driven by the constitutively active *Ubiquitin-10* promoter (Fuchs et al., 2020), was used to stably transform Col-0 and *mdh1xme1xme2.1* lines by floral dip (Clough and Bent, 1998). Two independent transgenic sensor lines (#1 and #2) were selected from the mutant background. For selection of primary transformants T0 seeds were germinated on 0.5x MS plates containing hygromycin B and resistant seedlings exhibiting mCherry fluorescence were identified using a Zeiss SteREO Discovery.V20 microscope with an HXP 120 C light source. For experiments seedlings from subsequent generations were cultivated without antibiotic but preselected using mCherry fluorescence.

For cytosolic NAD redox analysis, confocal imaging was performed using a Zeiss LSM 980 laser scanning microscope equipped with a 10x Plan-Apochromat objective (NA 0.3). The pinhole diameter was set to 145 µm. For fluorescence excitation and detection, the following settings were used: 405 nm excitation and 499-544 nm emission for tSapphire (Peredox), and 561 nm excitation and 588-624 nm emission for mCherry. Leaf discs used for imaging were excised from plants one hour before the onset of the light phase. Time-series acquisition was performed for 20 minutes in the dark to monitor the biosensor dynamics in the first abaxial mesophyll cell layer. Fluorescence data were processed using Redox Ratio Analysis Software (Fricker, 2016) with x-y noise filtering. For each line, at least three biological replicates were analyzed. From each replicate, the average of five representative time points was used.

### mRNA seq analysis

For RNA extraction, rosette leaves were harvested one hour before and after the light was switched on from plants growing in SD under LL and NL. Total RNA was extracted with the SV Total RNA Isolation System (Promega) and genomic DNA was eliminated by treating the samples with DNase I using DNA-free™ Kit (Invitrogen, Thermo Fischer), 1 µl of rDNase I was used per sample incubating at 37 °C for 30 min. The integrity of RNA was analyzed by gel electrophoresis, and the concentration was measured using a Nabi UV-Vis nano spectrophotometer (MicroDigital Co., Ltd., South Korea). Total RNA was quantified by the Qubit RNA HS Assay (Thermo Fisher Scientific, MA, USA). The quality was assessed by capillary electrophoresis using the Fragment Analyzer and the ‘Total RNA Standard Sensitivity Assay’ (Agilent Technologies, Inc. Santa Clara, CA, USA). Three biological replicates per genotype and condition with RNA Quality Numbers (RQN) mean > 8.2 were analyzed. The library preparation was performed according to the manufacturer’s protocol using the ‘VAHTS™ Stranded mRNA-Seq Library Prep Kit’ for Illumina®. Shortly, 400 ng total RNA were used as input for mRNA capturing, fragmentation, the synthesis of cDNA, adapter ligation and library amplification. Bead purified libraries were normalized and finally sequenced on the NextSeq2000 system (Illumina Inc. San Diego, CA, USA) with a read setup of 1x100 bp. The BCL Convert Tool (version 3.8.4) was used to convert the BCL files to FASTQ files as well for adapter trimming and demultiplexing.

Data analyses on FASTQ files were conducted with CLC Genomics Workbench (version 23.0.2, Qiagen, Venlo, Netherlands). The reads of all probes were adapter trimmed (Illumina TruSeq) and quality trimmed (using the default parameters: bases below Q13 were trimmed from the end of the reads, ambiguous nucleotides maximal 2). Mapping was done against the *A. thaliana* (TAIR10.54) (July 20, 2022) genome sequence. The differential expression analysis was performed using the DEseq2 package (Love et al., 2014) from BiocManager in the software RStudio, gene with less than five reads were depreciated. The Resulting *p* values calculated by Wald test were corrected for multiple testing using Benjamini and Hochberg method (BH-adjusted *p* values). An adjusted *p* value of ≤ 0.05 was considered significant.

### Shotgun proteomic LC-MS analysis

Proteomic analysis was performed on plants grown under SD conditions at light intensities of LL and NL. Whole rosettes were harvested 1.5 hours after the onset of light. Five biological replicates per genotype were collected. Protein extraction and digestion was carried out using a single-pot, solid-phase-enhanced sample preparation (SP3) protocol (Hughes et al., 2019), which was adapted to meet challenges inherent to plant samples (Mikulášek et al., 2021). Each extract was treated with 2 µg of pre-activated, sequencing grade Trypsin (V5111, Promega). Concentrations of extracted tryptic peptides were determined in a plate reader (Multiskan Sky, Thermo Fischer Scientific) using the Pierce Quantitative Colorimetric Peptide Assay Kit (Thermo Fisher Scientific, Dreieich, Germany) following the manufacturer’s instructions. Peptides were dried using a vacuum concentrator and resuspended in 0.1% (v/v) formic acid (FA) to yield a peptide concentration of 10 ng µl^-1^. Twenty microliters of the suspension were loaded on Evotips (Evosep) according to the manufacturer’s instructions for desalting and subsequent separation using an EvoSep One HPLC (Evosep). Peptides bound to the Evotip resin were eluted using the Whisper Zoom 40 SPD method and separated in a 15 cm Aurora Elite CSI column (inner diameter 75 µm, particle size 1.7 µm, pore size 120 Å, IonOpticks, Fitzroy, Australia) which was kept at a temperature of 50 °C. Eluting peptides were transferred to a timsTOF Pro mass spectrometer (Bruker, Germany) and analyzed using the pre-installed ‘DDA PASEF-short_gradient_0.5sec_cycletime’ method.

Acquired spectra were queried against an in-house modified TAIR10 database using the MaxQuant software (Cox and Mann, 2008) version 2.6.3.0. Default parameters were used, and the ‘iBAQ’ (Schwanhäusser et al., 2011) and ‘LFQ’ functions were additionally enabled for quantification of identified protein groups.

The resulting ‘proteingroups.txt’ file was complemented with predicted subcellular protein localization using the SUBAcon algorithm (Hooper et al., 2014) and Mapman annotation (Thimm et al., 2004). Clasified table was separated in two tables: plastid-protein groups and non-plastid-protein groups filtered by SUBAcon. These tables were submitted to statistical analysis by means of the Perseus software (Tyanova et al., 2016) version 2.1.2.0 in the form of two-sample t tests (*p* < 0.05). For this, protein groups matching ‘only identified by site’, ‘potential contaminant’ and ‘reverse’ were removed before LFQ values were log_2_-transformed. Protein groups with less than three values in each sample group were removed and missing values were imputed from normal distribution using a width of 0.3 and a down-shift of 1.8. A width normalization was performed volcano plots were produced using default parameters (FDR, 0.05; S_0_, 0.1).

### C/N ratio determination

For the determination of C/N ratio, whole rosettes were harvested one hour before the end of the night (-1), one hour after (+1) and seven hours after (+7) the light was switched on from plants growing under LL and under NL in SD. Four biological replicates were analyzed per genotype, 150 mg of fresh weight were dried in an oven at 60 °C for 72h, consequently samples were homogenized into a fine powder using a TissueLyserII (QIAGEN, Germany). Grinding was performed in two cycles of 1 minute each at a frequency of 30 Hz to ensure consistent pulverization. After that, 1.4 ± 0.2 mg of dry weight were placed into tin capsules (4x4x11 mm; Elementar Analysensysteme GmbH, Langenselbold). The quantification of total carbon and nitrogen was performed using Isoprime 100 isotope ratio mass spectrometer coupled to an ISOTOPE cube elemental analyzer (both from Elementar, Hanau, Germany) according to Schlüter et al. (Schlüter et al., 2023). For comparing the different genotypes, statistical analysis was done using). Results were statistically analyzed using a two-way ANOVA followed by Tukey’s post-hoc test using multcomp package (version:1.4-29; (Hothorn et al., 2008)) in R studio. *P*-val < 0.05 between the genotypes are considered significant differences.

### Ammonium concentration

For ammonium quantification whole rosette was harvested one hour before and after the light was on from plants growing under LL and NL in SD condition. The determination was done following a modified protocol from Vega-Mas, Sarasketa and Marino (Vega-Mas et al., 2015). Briefly, around 20 mg of frozen plant tissue was grinded using a TissueLyserII (QIAGEN, Germany) for 30 seconds two times at a frequency of 30 Hz, the third grinding step was done adding 1000 µl of HPLC quality Water. Following by an incubation for 10 min at 80°C and afterwards centrifugated 20 min at 9300 g at 4 °C. 50 µl of each sample was placed in a 96 well plate and the determination of free NH_4_^+^ was done adding to each well: 100 µl of 0.33 M sodium phenolate (Sigma-Aldrich), 50 µl of 0.02 % (w/v) sodium nitroprusside (Sigma-Aldrich), and 100 µl of Solution 2 % (v/v) sodium hypochlorite. The plate was incubated at room temperature for 30 min and the absorbance was measured at 635 nm. A calibration curve was run with (NH_4_)_2_SO_4_ in parallel. The value for each biological replicate was calculated as the average of three technical replicates. Results were statistically analyzed using a two-way ANOVA followed by Tukey’s post-hoc test using multcomp package (version:1.4-29; (Hothorn et al., 2008)) in R studio. *P*-val < 0.05 between the genotypes are considered significant differences.

### Metabolite profiling analysis

The metabolic profiles of the different lines were measured using gas chromatography–mass spectrometry (GC-MS). The whole rosette leaves were harvested at three time points, one hour before the light was on (-1), one hour after light was on (+1) and one hour before the light was off (+7/+15) from plants grown under SD-LL, SD-NL and LD-NL and immediately frozen. Approximately 50 mg of frozen tissue was ground two times for 30 seconds using a TissueLyserII (QIAGEN, Germany) at a frequency of 30 Hz. Soluble metabolites were extracted by a two phases extraction using six biological replicates for each line. After homogenization of the tissue, 700 µl of MeOH containing 30 ng/ µl Ribitol (Sigma-Aldrich) as standard compound was added followed by vertexing and shaking at 4 °C for 15 min. 700 µl of HPLC quality water and 350 µl of CHCl_3_ were added. Samples were centrifuged 4 min at 16,300 g for the phase separation. 300 µl of the polar phase were transferred to new tube and dried in a vacuum concentrator. Samples were subsequently derivatized for GC-MS analysis, as described by Gu et al (Gu et al., 2012). GC-MS measurements were carried out using a 5977B GC-MSD system (Agilent Technologies), following the procedure outlined by Shim et al (Shim et al., 2020). Metabolites were identified using MassHunter Qualitative Analysis software (version B.08.00, Agilent Technologies) by comparing acquired spectra with entries in the NIST14 Mass Spectral Library (https://www.nist.gov/srd/nist-standard-reference-database-1a-v14). A standard mixture containing all target compounds at a concentration of 5 µM was processed alongside the samples to serve as quality control for response consistency and retention time referencing. Peak integration was performed using MassHunter Quantitative Analysis software (version B.08.00, Agilent Technologies). For relative quantification, metabolite peak areas were normalized to the fresh weight of the sample and to the peak area of the internal standard (Ribitol, Sigma-Aldrich). Results were statistically analyzed using a two-way ANOVA followed by Tukey’s post-hoc test, multcomp (version:1.4-29; (Hothorn et al., 2008)) package was used in R studio. Using a threshold of *p* ≤ 0.05 between the samples of the control and mutant plants (Supplementary Data 3).

## Data availability

The data supporting the findings of this manuscript are available from the corresponding author upon reasonable request. The sequencing data generated for this study have been deposited in the European Nucleotide Archive (ENA) under accession number PRJEB96949 and are publicly available at PRJEB96949 (https://www.ebi.ac.uk/ena/browser/view/PRJEB96949). The raw RNA-seq data (FASTQ files) supporting the findings of this study are available in the NCBI SRA repository, under BioProject accession PRJNA1427118 (https://www.ncbi.nlm.nih.gov/sra/PRJNA1427118). All MS raw files acquired in this study, as well as the corresponding search engine results produced by MaxQuant can be downloaded under ftp://MSV000100916@massive-ftp.ucsd.edu. Reviewer login: MSV000100916_reviewer; password: MalateMetabolism, or on the MassIve website using the above-mentioned login data.

## Author contributions

V.M. and M.P.M conceived the study. M.P.M., I.N., K.Z., N.D., J.A.V.S.O., P.B., N.B., P.W., and H.E. performed experiments and collected data. V.M., M.P.M., H.E., I.F. and M.S. analyzed the data. V.M., M.P.M., I.F., and M.S. interpreted results. V.G.M and M.P.M. wrote the manuscript with input from all authors. V.M. supervised the project. V.M., I.F., and M.S. acquired funding.

## Acknowledgements

We acknowledge support from the DFG collaborative research grant PAK918 (Project ID 289357231; MA2379/14-3; FI 1655/3-3; SCHW 1719/5-3) within the “Plant Mitochondria in New Light” initiative. We thank the CEPLAS Metabolism & Metabolomics Laboratory (CMML; Heinrich Heine University Düsseldorf) for technical assistance; CMML is supported by the DFG (CEPLAS, EXC-2048/1; Project ID 390686111). Computational infrastructure and support for RNA-seq analyses were provided by the Centre for Information and Media Technology at Heinrich Heine University Düsseldorf, with additional support from the de.NBI Cloud within the German Network for Bioinformatics Infrastructure (de.NBI) and ELIXIR-DE (Forschungszentrum Jülich and W-de.NBI-001, W-de.NBI-004, W-de.NBI-008, W-de.NBI-010, W-de.NBI-013, W-de.NBI-014, W-de.NBI-016, W-de.NBI-022).

The co-authors have no conflict of interest to declare.

## REFERENCES

1. Albanese P, Tamara S, Saracco G, Scheltema RA, Pagliano C (2020) How paired PSII–LHCII supercomplexes mediate the stacking of plant thylakoid membranes unveiled by structural mass-spectrometry. Nat Commun 11: 1361

2. Araújo WL, Ishizaki K, Nunes-Nesi A, Larson TR, Tohge T, Krahnert I, Witt S, Obata T, Schauer N, Graham IA, et al (2010) Identification of the 2-Hydroxyglutarate and Isovaleryl-CoA Dehydrogenases as Alternative Electron Donors Linking Lysine Catabolism to the Electron Transport Chain of Arabidopsis Mitochondria. Plant Cell 22: 1549–1563

3. Arioli T, Peng L, Betzner AS, Burn J, Wittke W, Herth W, Camilleri C, Höfte H, Plazinski J, Birch R, et al (1998) Molecular Analysis of Cellulose Biosynthesis in Arabidopsis. Science 279: 717–720

4. Awai K, Maréchal E, Block MA, Brun D, Masuda T, Shimada H, Takamiya K, Ohta H, Joyard J (2001) Two types of MGDG synthase genes, found widely in both 16:3 and 18:3 plants, differentially mediate galactolipid syntheses in photosynthetic and nonphotosynthetic tissues in Arabidopsis thaliana. Proceedings of the National Academy of Sciences 98: 10960–10965

5. Bachmann M, Matile P, Keller F (1994) Metabolism of the Raffinose Family Oligosaccharides in Leaves of Ajuga reptans L. (Cold Acclimation, Translocation, and Sink to Source Transition: Discovery of Chain Elongation Enzyme). Plant Physiol 105: 1335–1345

6. Balparda M, Elsässer M, Badia MB, Giese J, Bovdilova A, Hüdig M, Reinmuth L, Eirich J, Schwarzländer M, Finkemeier I, et al (2022) Acetylation of conserved lysines fine-tunes mitochondrial malate dehydrogenase activity in land plants. The Plant Journal 109: 92–111

7. Basso L, Yamori W, Szabo I, Shikanai T (2020) Collaboration between NDH and KEA3 Allows Maximally Efficient Photosynthesis after a Long Dark Adaptation. Plant Physiol 184: 2078–2090

8. Ben-Shem A, Frolow F, Nelson N (2003) Crystal structure of plant photosystem I. Nature 426: 630–635

9. Brauc S, De Vooght E, Claeys M, Geuns JMC, Höfte M, Angenon G (2012) Overexpression of arginase in Arabidopsis thaliana influences defence responses against Botrytis cinerea. Plant Biology 14: 39–45

10. Briat J-F, Lobréaux S (1997) Iron transport and storage in plants. Trends in Plant Science 2: 187–193

11. Bykova NV, Møller IM, Gardeström P, Igamberdiev AU (2014) The function of glycine decarboxylase complex is optimized to maintain high photorespiratory flux via buffering of its reaction products. Mitochondrion 19 Pt B: 357–364

12. Camp PJ, Randall DD (1985) Purification and Characterization of the Pea Chloroplast Pyruvate Dehydrogenase Complex 1: A Source of Acetyl-CoA and NADH for Fatty Acid Biosynthesis. Plant Physiol 77: 571–577

13. Carmi N, Zhang G, Petreikov M, Gao Z, Eyal Y, Granot D, Schaffer AA (2003) Cloning and functional expression of alkaline α-galactosidase from melon fruit: similarity to plant SIP proteins uncovers a novel family of plant glycosyl hydrolases. The Plant Journal 33: 97–106

14. Che P, Wurtele ES, Nikolau BJ (2002) Metabolic and Environmental Regulation of 3-Methylcrotonyl-Coenzyme A Carboxylase Expression in Arabidopsis. Plant Physiol 129: 625–637

15. Clough SJ, Bent AF (1998) Floral dip: a simplified method for Agrobacterium -mediated transformation of Arabidopsis thaliana. The Plant Journal 16: 735–743

16. Cox J, Mann M (2008) MaxQuant enables high peptide identification rates, individualized p.p.b.-range mass accuracies and proteome-wide protein quantification. Nat Biotechnol 26: 1367–1372

17. De Vylder J, Vandenbussche F, Hu Y, Philips W, Van Der Straeten D (2012) Rosette Tracker: An Open Source Image Analysis Tool for Automatic Quantification of Genotype Effects. Plant Physiol 160: 1149–1159

18. Diebold R, Schuster J, Däschner K, Binder S (2002) The Branched-Chain Amino Acid Transaminase Gene Family in Arabidopsis Encodes Plastid and Mitochondrial Proteins. Plant Physiol 129: 540–550

19. Dieuaide-Noubhani M, Raffard G, Canioni P, Pradet A, Raymond P (1995) Quantification of compartmented metabolic fluxes in maize root tips using isotope distribution from 13C- or 14C-labeled glucose. J Biol Chem 270: 13147–13159

20. Ding G, Che P, Ilarslan H, Wurtele ES, Nikolau BJ (2012) Genetic dissection of methylcrotonyl CoA carboxylase indicates a complex role for mitochondrial leucine catabolism during seed development and germination. The Plant Journal 70: 562–577

21. Drath M, Kloft N, Batschauer A, Marin K, Novak J, Forchhammer K (2008) Ammonia triggers photodamage of photosystem II in the cyanobacterium Synechocystis sp. strain PCC 6803. Plant Physiol 147: 206–215

22. Drincovich MF, Voll LM, Maurino VG (2016) Editorial: On the Diversity of Roles of Organic Acids. Front Plant Sci 7: 1592

23. Engqvist MKM, Kuhn A, Wienstroer J, Weber K, Jansen EEW, Jakobs C, Weber APM, Maurino VG (2011) Plant d-2-Hydroxyglutarate Dehydrogenase Participates in the Catabolism of Lysine Especially during Senescence. Journal of Biological Chemistry 286: 11382–11390

24. Farber A, Young AJ, Ruban AV, Horton P, Jahns P (1997) Dynamics of Xanthophyll-Cycle Activity in Different Antenna Subcomplexes in the Photosynthetic Membranes of Higher Plants (The Relationship between Zeaxanthin Conversion and Nonphotochemical Fluorescence Quenching). Plant Physiol 115: 1609–1618

25. Fontaine J-X, Tercé-Laforgue T, Armengaud P, Clément G, Renou J-P, Pelletier S, Catterou M, Azzopardi M, Gibon Y, Lea PJ, et al (2012) Characterization of a NADH-Dependent Glutamate Dehydrogenase Mutant of Arabidopsis Demonstrates the Key Role of this Enzyme in Root Carbon and Nitrogen Metabolism. Plant Cell 24: 4044–4065

26. Fricker MD (2016) Quantitative Redox Imaging Software. Antioxidants & Redox Signaling 24: 752–762

27. Fuchs P, Rugen N, Carrie C, Elsässer M, Finkemeier I, Giese J, Hildebrandt TM, Kühn K, Maurino VG, Ruberti C, et al (2020) Single organelle function and organization as estimated from Arabidopsis mitochondrial proteomics. Plant J 101: 420–441

28. Fujiki Y, Yoshikawa Y, Sato T, Inada N, Ito M, Nishida I, Watanabe A (2001) Dark-inducible genes from Arabidopsis thaliana are associated with leaf senescence and repressed by sugars. Physiologia Plantarum 111: 345–352

29. Gaudreault P-R, Webb JA (1986) Alkaline α-galactosidase activity and galactose metabolism in the family *Cucurbitaceae*. Plant Science 45: 71–75

30. Giese J, Eirich J, Walther D, Zhang Y, Lassowskat I, Fernie AR, Elsässer M, Maurino VG, Schwarzländer M, Finkemeier I (2023) The interplay of post-translational protein modifications in Arabidopsis leaves during photosynthesis induction. The Plant Journal 116: 1172–1193

31. Gietl C (1992) Partitioning of malate dehydrogenase isoenzymes into glyoxysomes, mitochondria, and chloroplasts. Plant Physiol 100: 557–559

32. Gu J, Weber K, Klemp E, Winters G, Franssen SU, Wienpahl I, Huylmans A-K, Zecher K, Reusch TBH, Bornberg-Bauer E, et al (2012) Identifying core features of adaptive metabolic mechanisms for chronic heat stress attenuation contributing to systems robustness. Integr Biol 4: 480–493

33. Hachiya T, Inaba J, Wakazaki M, Sato M, Toyooka K, Miyagi A, Kawai-Yamada M, Sugiura D, Nakagawa T, Kiba T, et al (2021) Excessive ammonium assimilation by plastidic glutamine synthetase causes ammonium toxicity in Arabidopsis thaliana. Nat Commun 12: 4944

34. Hauser T, Bhat JY, Miličić G, Wendler P, Hartl FU, Bracher A, Hayer-Hartl M (2015) Structure and mechanism of the Rubisco-assembly chaperone Raf1. Nat Struct Mol Biol 22: 720–728

35. Herrmann HA, Dyson BC, Miller MAE, Schwartz J-M, Johnson GN (2021) Metabolic flux from the chloroplast provides signals controlling photosynthetic acclimation to cold in Arabidopsis thaliana. Plant, Cell & Environment 44: 171–185

36. Hooper CM, Tanz SK, Castleden IR, Vacher MA, Small ID, Millar AH (2014) SUBAcon: a consensus algorithm for unifying the subcellular localization data of the Arabidopsis proteome. Bioinformatics 30: 3356–3364

37. Hothorn T, Bretz F, Westfall P (2008) Simultaneous Inference in General Parametric Models. Biometrical Journal 50: 346–363

38. Hüdig M, Maier A, Scherrers I, Seidel L, Jansen EEW, Mettler-Altmann T, Engqvist MKM, Maurino VG (2015) Plants Possess a Cyclic Mitochondrial Metabolic Pathway similar to the Mammalian Metabolic Repair Mechanism Involving Malate Dehydrogenase and l-2-Hydroxyglutarate Dehydrogenase. Plant Cell Physiol 56: 1820–1830

39. Hughes RA, Heron J, Sterne JAC, Tilling K (2019) Accounting for missing data in statistical analyses: multiple imputation is not always the answer. Int J Epidemiol 48: 1294–1304

40. Ishihara S, Takabayashi A, Ido K, Endo T, Ifuku K, Sato F (2007) Distinct Functions for the Two PsbP-Like Proteins PPL1 and PPL2 in the Chloroplast Thylakoid Lumen of Arabidopsis. Plant Physiol 145: 668–679

41. Ishizaki K, Larson TR, Schauer N, Fernie AR, Graham IA, Leaver CJ (2005) The Critical Role of Arabidopsis Electron-Transfer Flavoprotein:Ubiquinone Oxidoreductase during Dark-Induced Starvation. Plant Cell 17: 2587–2600

42. Izumi M, Tsunoda H, Suzuki Y, Makino A, Ishida H (2012) RBCS1A and RBCS3B, two major members within the Arabidopsis RBCS multigene family, function to yield sufficient Rubisco content for leaf photosynthetic capacity. J Exp Bot 63: 2159–2170

43. Jiang T, Du K, Xie J, Sun G, Wang P, Chen X, Cao Z, Wang B, Chao Q, Li X, et al (2023) Activated malate circulation contributes to the manifestation of light-dependent mosaic symptoms. Cell Reports 42: 112333

44. Johnson MP, Ruban AV (2011) Restoration of Rapidly Reversible Photoprotective Energy Dissipation in the Absence of PsbS Protein by Enhanced ΔpH*. Journal of Biological Chemistry 286: 19973–19981

45. Ke J, Behal RH, Back SL, Nikolau BJ, Wurtele ES, Oliver DJ (2000) The role of pyruvate dehydrogenase and acetyl-coenzyme A synthetase in fatty acid synthesis in developing Arabidopsis seeds. Plant Physiol 123: 497–508

46. Kleinboelting N, Huep G, Kloetgen A, Viehoever P, Weisshaar B (2012) GABI-Kat SimpleSearch: new features of the Arabidopsis thaliana T-DNA mutant database. Nucleic Acids Res 40: D1211–D1215

47. Kobayashi K, Awai K, Nakamura M, Nagatani A, Masuda T, Ohta H (2009) Type-B monogalactosyldiacylglycerol synthases are involved in phosphate starvation-induced lipid remodeling, and are crucial for low-phosphate adaptation. The Plant Journal 57: 322–331

48. Kobayashi K, Awai K, Takamiya K, Ohta H (2004) Arabidopsis Type B Monogalactosyldiacylglycerol Synthase Genes Are Expressed during Pollen Tube Growth and Induced by Phosphate Starvation. Plant Physiol 134: 640–648

49. Kochian LV (1995) Cellular Mechanisms of Aluminum Toxicity and Resistance in Plants. Annu Rev Plant Physiol Plant Mol Biol 46: 237–260

50. Kou J, Takahashi S, Fan D-Y, Badger MR, Chow WS (2015) Partially dissecting the steady-state electron fluxes in Photosystem I in wild-type and pgr5 and ndh mutants of Arabidopsis. Front Plant Sci 6: 758

51. Kouno T, Ezaki B (2013) Multiple regulation of Arabidopsis AtGST11 gene expression by four transcription factors under abiotic stresses. Physiologia Plantarum 148: 97–104

52. Kromer S (1995) Respiration During Photosynthesis. Annual Review of Plant Biology 46: 45–70

53. Krtková J, Benáková M, Schwarzerová K (2016) Multifunctional Microtubule-Associated Proteins in Plants. Front Plant Sci 7: 474

54. Lam HM, Peng SSY, Coruzzi GM (1994) Metabolic Regulation of the Gene Encoding Glutamine-Dependent Asparagine Synthetase in Arabidopsis thaliana. Plant Physiol 106: 1347–1357

55. Larbi A, Abadía A, Morales F, Abadía J (2004) Fe Resupply to Fe-deficient Sugar Beet Plants Leads to Rapid Changes in the Violaxanthin Cycle and other Photosynthetic Characteristics without Significant de novo Chlorophyll Synthesis. Photosynthesis Research 79: 59–69

56. Le XH, Lee CP, Monachello D, Millar AH (2022) Metabolic evidence for distinct pyruvate pools inside plant mitochondria. Nat Plants 8: 694–705

57. Lee CP, Eubel H, O’Toole N, Millar AH (2008) Heterogeneity of the mitochondrial proteome for photosynthetic and non-photosynthetic Arabidopsis metabolism. Mol Cell Proteomics 7: 1297–1316

58. Lee R-H, Hsu J-H, Huang H-J, Lo S-F, Grace Chen S-C (2009) Alkaline α-galactosidase degrades thylakoid membranes in the chloroplast during leaf senescence in rice. New Phytologist 184: 596–606

59. Li H (2018) Minimap2: pairwise alignment for nucleotide sequences. Bioinformatics 34: 3094–3100

60. Li H, Handsaker B, Wysoker A, Fennell T, Ruan J, Homer N, Marth G, Abecasis G, Durbin R, 1000 Genome Project Data Processing Subgroup (2009) The Sequence Alignment/Map format and SAMtools. Bioinformatics 25: 2078–2079

61. Li J, Xu Y, Chong K (2012) The novel functions of kinesin motor proteins in plants. Protoplasma 249: 95–100

62. Li Q, Hoppe T (2023) Role of amino acid metabolism in mitochondrial homeostasis. Front Cell Dev Biol 11: 1127618

63. Li X-P, Björkman O, Shih C, Grossman AR, Rosenquist M, Jansson S, Niyogi KK (2000) A pigment-binding protein essential for regulation of photosynthetic light harvesting. Nature 403: 391–395

64. Lin J-F, Wu S-H (2004) Molecular events in senescing Arabidopsis leaves. The Plant Journal 39: 612–628

65. Liu J, Last RL (2017) A chloroplast thylakoid lumen protein is required for proper photosynthetic acclimation of plants under fluctuating light environments. Proceedings of the National Academy of Sciences 114: E8110–E8117

66. Lobo AKM, Orr DJ, Gutierrez MO, Andralojc PJ, Sparks C, Parry MAJ, Carmo-Silva E (2019) Overexpression of ca1pase Decreases Rubisco Abundance and Grain Yield in Wheat1 [CC-BY]. Plant Physiol 181: 471–479

67. Love MI, Huber W, Anders S (2014) Moderated estimation of fold change and dispersion for RNA-seq data with DESeq2. Genome Biol 15: 550

68. Martín M, Noarbe DM, Serrot PH, Sabater B (2015) The rise of the photosynthetic rate when light intensity increases is delayed in ndh gene-defective tobacco at high but not at low CO2 concentrations. Front Plant Sci 6: 34

69. Martinoia E, Rentsch D (1994) Malate Compartmentation-Responses to a Complex Metabolism. Annu Rev Plant Physiol Plant Mol Biol 45: 447–467

70. Maurino VG, Engqvist MKM (2015) 2-Hydroxy Acids in Plant Metabolism. The Arabidopsis Book 13: e0182

71. May A, Berger S, Hertel T, Köck M (2011) The *Arabidopsis thaliana* phosphate starvation responsive gene *AtPPsPase1* encodes a novel type of inorganic pyrophosphatase. Biochimica et Biophysica Acta (BBA) - General Subjects 1810: 178–185

72. McElver J, Tzafrir I, Aux G, Rogers R, Ashby C, Smith K, Thomas C, Schetter A, Zhou Q, Cushman MA, et al (2001) Insertional Mutagenesis of Genes Required for Seed Development in Arabidopsis thaliana. Genetics 159: 1751–1763

73. Mikulášek K, Konečná H, Potěšil D, Holánková R, Havliš J, Zdráhal Z (2021) SP3 Protocol for Proteomic Plant Sample Preparation Prior LC-MS/MS. Front Plant Sci 12: 635550

74. Miyashita Y, Good AG (2008) NAD(H)-dependent glutamate dehydrogenase is essential for the survival of Arabidopsis thaliana during dark-induced carbon starvation. J Exp Bot 59: 667–680

75. Møller IM, Rasmusson AG (1998) The role of NADP in the mitochondrial matrix. Trends in Plant Science 3: 21–27

76. Moreno-García B, López-Calcagno PE, Raines CA, Sweetlove LJ (2022) Suppression of metabolite shuttles for export of chloroplast and mitochondrial ATP and NADPH increases the cytosolic NADH:NAD+ ratio in tobacco leaves in the dark. Journal of Plant Physiology 268: 153578

77. Murgia I, Vazzola V, Tarantino D, Cellier F, Ravet K, Briat J-F, Soave C (2007) Knock-out of ferritin *AtFer1* causes earlier onset of age-dependent leaf senescence in *Arabidopsis*. Plant Physiology and Biochemistry 45: 898–907

78. Nakano Y, Naito Y, Nakano T, Ohtsuki N, Suzuki K (2017) *NSR1/MYR2* is a negative regulator of *ASN1* expression and its possible involvement in regulation of nitrogen reutilization in Arabidopsis. Plant Science 263: 219–225

79. Netting AG (2000) pH, abscisic acid and the integration of metabolism in plants under stressed and non-stressed conditions: cellular responses to stress and their implication for plant water relations. J Exp Bot 51: 147–158

80. Nozawa A, Ito M, Hayashi H, Watanabe A (1999) Dark-Induced Expression of Genes for Asparagine Synthetase and Cytosolic Glutamine Synthetase in Radish Cotyledons is Dependent on the Growth Stage. Plant and Cell Physiology 40: 942–948

81. de Oliveira Dal’Molin CG, Quek L-E, Palfreyman RW, Brumbley SM, Nielsen LK (2010) AraGEM, a Genome-Scale Reconstruction of the Primary Metabolic Network in Arabidopsis. Plant Physiol 152: 579–589

82. Palmieri F, Rieder B, Ventrella A, Blanco E, Do PT, Nunes-Nesi A, Trauth AU, Fiermonte G, Tjaden J, Agrimi G, et al (2009) Molecular identification and functional characterization of Arabidopsis thaliana mitochondrial and chloroplastic NAD+ carrier proteins. J Biol Chem 284: 31249–31259

83. Palmieri L, Santoro A, Carrari F, Blanco E, Nunes-Nesi A, Arrigoni R, Genchi F, Fernie AR, Palmieri F (2008) Identification and Characterization of ADNT1, a Novel Mitochondrial Adenine Nucleotide Transporter from Arabidopsis. Plant Physiol 148: 1797–1808

84. Peng C, Uygun S, Shiu S-H, Last RL (2015) The Impact of the Branched-Chain Ketoacid Dehydrogenase Complex on Amino Acid Homeostasis in Arabidopsis. Plant Physiol 169: 1807–1820

85. Persson S, Paredez A, Carroll A, Palsdottir H, Doblin M, Poindexter P, Khitrov N, Auer M, Somerville CR (2007) Genetic evidence for three unique components in primary cell-wall cellulose synthase complexes in Arabidopsis. Proceedings of the National Academy of Sciences 104: 15566–15571

86. Peterson RB (2005) PsbS genotype in relation to coordinated function of PS II and PS I in Arabidopsis leaves. Photosynth Res 85: 205–219

87. Popov VN, Eprintsev AT, Fedorin DN, Igamberdiev AU (2010) Succinate dehydrogenase in Arabidopsis thaliana is regulated by light via phytochrome A. FEBS Letters 584: 199–202

88. Post-Beittenmiller D, Roughan G, Ohlrogge JB (1992) Regulation of plant Fatty Acid biosynthesis : analysis of acyl-coenzyme a and acyl-acyl carrier protein substrate pools in spinach and pea chloroplasts. Plant Physiol 100: 923–930

89. Pribil M, Pesaresi P, Hertle A, Barbato R, Leister D (2010) Role of Plastid Protein Phosphatase TAP38 in LHCII Dephosphorylation and Thylakoid Electron Flow. PLOS Biology 8: e1000288

90. Pu MN, Liang G (2023) The transcription factor POPEYE negatively regulates the expression of bHLH Ib genes to maintain iron homeostasis. J Exp Bot 74: 2754–2767

91. Pucker B, Brockington SF (2018) Genome-wide analyses supported by RNA-Seq reveal non-canonical splice sites in plant genomes. BMC Genomics 19: 980

92. Pucker B, Kleinbölting N, Weisshaar B (2021) Large scale genomic rearrangements in selected Arabidopsis thaliana T-DNA lines are caused by T-DNA insertion mutagenesis. BMC Genomics 22: 599

93. Quinlan AR, Hall IM (2010) BEDTools: a flexible suite of utilities for comparing genomic features. Bioinformatics 26: 841–842

94. Raghavendra AS, Padmasree K (2003) Beneficial interactions of mitochondrial metabolism with photosynthetic carbon assimilation. Trends Plant Sci 8: 546–553

95. Rasmusson AG, Escobar MA (2007) Light and diurnal regulation of plant respiratory gene expression. Physiologia Plantarum 129: 57–67

96. Ravet K, Touraine B, Boucherez J, Briat J-F, Gaymard F, Cellier F (2009) Ferritins control interaction between iron homeostasis and oxidative stress in Arabidopsis. The Plant Journal 57: 400–412

97. Reiter W-D, Vanzin GF (2001) Molecular genetics of nucleotide sugar interconversion pathways in plants. Plant Mol Biol 47: 95–113

98. Robinson JT, Thorvaldsdóttir H, Winckler W, Guttman M, Lander ES, Getz G, Mesirov JP (2011) Integrative genomics viewer. Nat Biotechnol 29: 24–26

99. Ryan PR, Delhaize E, Jones DL (2001) Function and mechanism of organic anion exudation from pant roots. Annual Review of Plant Biology 52: 527–560

100. Santelia D, Lawson T (2016) Rethinking Guard Cell Metabolism. Plant Physiol 172: 1371–1392

101. Sappl PG, Carroll AJ, Clifton R, Lister R, Whelan J, Millar AH, Singh KB (2009) The Arabidopsis glutathione transferase gene family displays complex stress regulation and co-silencing multiple genes results in altered metabolic sensitivity to oxidative stress. The Plant Journal 58: 53–68

102. Schachtman DP, Reid RJ, Ayling SM (1998) Phosphorus Uptake by Plants: From Soil to Cell. Plant Physiol 116: 447–453

103. Scheibe R (2004) Malate valves to balance cellular energy supply. Physiol Plant 120: 21–26

104. Schlüter U, Bouvier JW, Guerreiro R, Malisic M, Kontny C, Westhoff P, Stich B, Weber APM (2023) *Brassicaceae* display variation in efficiency of photorespiratory carbon-recapturing mechanisms. J Exp Bot 74: 6631–6649

105. Schuster J, Binder S (2005) The mitochondrial branched-chain aminotransferase (AtBCAT-1) is capable to initiate degradation of leucine, isoleucine and valine in almost all tissues in *Arabidopsis thaliana*. Plant Mol Biol 57: 241–254

106. Schwanhäusser B, Busse D, Li N, Dittmar G, Schuchhardt J, Wolf J, Chen W, Selbach M (2011) Global quantification of mammalian gene expression control. Nature 473: 337–342

107. Seifert GJ (2018) Mad moves of the building blocks – nucleotide sugars find unexpected paths into cell walls. J Exp Bot 69: 905–907

108. Selinski J, Scheibe R (2019) Malate valves: old shuttles with new perspectives. Plant Biol (Stuttg) 21 Suppl 1: 21–30

109. Sew YS, Ströher E, Fenske R, Millar AH (2016) Loss of Mitochondrial Malate Dehydrogenase Activity Alters Seed Metabolism Impairing Seed Maturation and Post-Germination Growth in Arabidopsis. Plant Physiol 171: 849–863

110. Shameer S, Ratcliffe RG, Sweetlove LJ (2019) Leaf Energy Balance Requires Mitochondrial Respiration and Export of Chloroplast NADPH in the Light. Plant Physiol 180: 1947–1961

111. Shapiguzov A, Ingelsson B, Samol I, Andres C, Kessler F, Rochaix J-D, Vener AV, Goldschmidt-Clermont M (2010) The PPH1 phosphatase is specifically involved in LHCII dephosphorylation and state transitions in Arabidopsis. Proceedings of the National Academy of Sciences 107: 4782–4787

112. Shim S-H, Lee S-K, Lee D-W, Brilhaus D, Wu G, Ko S, Lee C-H, Weber APM, Jeon J-S (2020) Loss of Function of Rice Plastidic Glycolate/Glycerate Translocator 1 Impairs Photorespiration and Plant Growth. Front Plant Sci 10: 1726

113. Siadjeu C, Pucker B, Viehöver P, Albach DC, Weisshaar B (2020) High Contiguity *de novo* Genome Sequence Assembly of Trifoliate Yam (*Dioscorea dumetorum*) Using Long Read Sequencing. Genes (Basel) 11: 274

114. Spiller S, Terry N (1980) Limiting Factors in Photosynthesis: II. iron stress diminishes photochemical capacity by reducing the number of photosynthetic units. Plant Physiol 65: 121–125

115. Steinbeck J, Fuchs P, Negroni YL, Elsässer M, Lichtenauer S, Stockdreher Y, Feitosa-Araujo E, Kroll JB, Niemeier J-O, Humberg C, et al (2020) In Vivo NADH/NAD+ Biosensing Reveals the Dynamics of Cytosolic Redox Metabolism in Plants. Plant Cell 32: 3324–3345

116. Steuer R, Nesi AN, Fernie AR, Gross T, Blasius B, Selbig J (2007) From structure to dynamics of metabolic pathways: application to the plant mitochondrial TCA cycle. Bioinformatics 23: 1378–1385

117. Su T, Xu J, Li Y, Lei L, Zhao L, Yang H, Feng J, Liu G, Ren D (2011) Glutathione-Indole-3-Acetonitrile Is Required for Camalexin Biosynthesis in *Arabidopsis thaliana*. Plant Cell 23: 364–380

118. Sweetlove LJ, Beard KFM, Nunes-Nesi A, Fernie AR, Ratcliffe RG (2010) Not just a circle: flux modes in the plant TCA cycle. Trends Plant Sci 15: 462–470

119. Szklarczyk D, Kirsch R, Koutrouli M, Nastou K, Mehryary F, Hachilif R, Gable AL, Fang T, Doncheva NT, Pyysalo S, et al (2023) The STRING database in 2023: protein–protein association networks and functional enrichment analyses for any sequenced genome of interest. Nucleic Acids Res 51: D638–D646

120. Tcherkez G, Cornic G, Bligny R, Gout E, Ghashghaie J (2005) In vivo respiratory metabolism of illuminated leaves. Plant Physiol 138: 1596–1606

121. Thimm O, Bläsing O, Gibon Y, Nagel A, Meyer S, Krüger P, Selbig J, Müller LA, Rhee SY, Stitt M (2004) mapman: a user-driven tool to display genomics data sets onto diagrams of metabolic pathways and other biological processes. The Plant Journal 37: 914–939

122. Thum KE, Shin MJ, Palenchar PM, Kouranov A, Coruzzi GM (2004) Genome-wide investigation of light and carbon signaling interactions in Arabidopsis. Genome Biol 5: R10

123. Tomaz T, Bagard M, Pracharoenwattana I, Lindén P, Lee CP, Carroll AJ, Ströher E, Smith SM, Gardeström P, Millar AH (2010) Mitochondrial malate dehydrogenase lowers leaf respiration and alters photorespiration and plant growth in Arabidopsis. Plant Physiol 154: 1143–1157

124. Tronconi MA, Fahnenstich H, Gerrard Weehler MC, Andreo CS, Flügge U-I, Drincovich MF, Maurino VG (2008) Arabidopsis NAD-Malic Enzyme Functions As a Homodimer and Heterodimer and Has a Major Impact on Nocturnal Metabolism. Plant Physiol 146: 1540–1552

125. Tronconi MA, Maurino VG, Andreo CS, Drincovich MF (2010) Three Different and Tissue-specific NAD-Malic Enzymes Generated by Alternative Subunit Association in *Arabidopsis thaliana*. Journal of Biological Chemistry 285: 11870–11879

126. Tyanova S, Temu T, Cox J (2016) The MaxQuant computational platform for mass spectrometry-based shotgun proteomics. Nat Protoc 11: 2301–2319

127. Umena Y, Kawakami K, Shen J-R, Kamiya N (2011) Crystal structure of oxygen-evolving photosystem II at a resolution of 1.9 Å. Nature 473: 55–60

128. Vega-Mas I, Sarasketa A, Marino D (2015) High-throughput Quantification of Ammonium Content in Arabidopsis. Bio-protoco 5: e1559

129. Wagner S, Steinbeck J, Fuchs P, Lichtenauer S, Elsässer M, Schippers JHM, Nietzel T, Ruberti C, Van Aken O, Meyer AJ, et al (2019) Multiparametric real-time sensing of cytosolic physiology links hypoxia responses to mitochondrial electron transport. New Phytologist 224: 1668–1684

130. Wang N, Cui Y, Liu Y, Fan H, Du J, Huang Z, Yuan Y, Wu H, Ling H-Q (2013) Requirement and Functional Redundancy of Ib Subgroup bHLH Proteins for Iron Deficiency Responses and Uptake in *Arabidopsis thaliana*. Molecular Plant 6: 503–513

131. Whitney SM, Birch R, Kelso C, Beck JL, Kapralov MV (2015) Improving recombinant Rubisco biogenesis, plant photosynthesis and growth by coexpressing its ancillary RAF1 chaperone. Proceedings of the National Academy of Sciences 112: 3564–3569

132. Wilkinson JQ, Crawford NM (1993) Identification and characterization of a chlorate-resistant mutant of Arabidopsis thaliana with mutations in both nitrate reductase structural genes NIA1 and NIA2. Molec Gen Genet 239: 289–297

133. Williams TCR, Miguet L, Masakapalli SK, Kruger NJ, Sweetlove LJ, Ratcliffe RG (2008) Metabolic network fluxes in heterotrophic Arabidopsis cells: stability of the flux distribution under different oxygenation conditions. Plant Physiol 148: 704–718

134. Wu J, Sun Y, Zhao Y, Zhang J, Luo L, Li M, Wang J, Yu H, Liu G, Yang L, et al (2015) Deficient plastidic fatty acid synthesis triggers cell death by modulating mitochondrial reactive oxygen species. Cell Res 25: 621–633

135. Yamori W, Sakata N, Suzuki Y, Shikanai T, Makino A (2011) Cyclic electron flow around photosystem I via chloroplast NAD(P)H dehydrogenase (NDH) complex performs a significant physiological role during photosynthesis and plant growth at low temperature in rice. The Plant Journal 68: 966–976

136. Yoshida K, Hisabori T (2016) Two distinct redox cascades cooperatively regulate chloroplast functions and sustain plant viability. Proc Natl Acad Sci U S A 113: E3967–3976

137. Yuan Y, Wu H, Wang N, Li J, Zhao W, Du J, Wang D, Ling H-Q (2008) FIT interacts with AtbHLH38 and AtbHLH39 in regulating iron uptake gene expression for iron homeostasis in Arabidopsis. Cell Res 18: 385–397

138. Zhao Y, Luo L, Xu J, Xin P, Guo H, Wu J, Bai L, Wang G, Chu J, Zuo J, et al (2018) Malate transported from chloroplast to mitochondrion triggers production of ROS and PCD in Arabidopsis thaliana. Cell Res 28: 448–461

139. Zhao Y, Yu H, Zhou J-M, Smith SM, Li J (2020) Malate Circulation: Linking Chloroplast Metabolism to Mitochondrial ROS. Trends in Plant Science 25: 446–454

140. Zheng K, Elsässer M, Niemeier J-O, Barreto P, Cislaghi AP, Hoang M, Feitosa-Araujo E, Wagner S, Giese J, Kotnik F, et al (2025) Advanced illumination-imaging reveals photosynthesis-triggered pH, ATP and NAD redox signatures across plant cell compartments. doi: 10.1101/2025.06.16.659786

