## Supplementary Figures and Tables for "Mitochondrial malate metabolism acts as a control hub for photosynthesis and carbon-nitrogen balance in Arabidopsis"

### Supplementary Figures (Martinez et al., 2026)

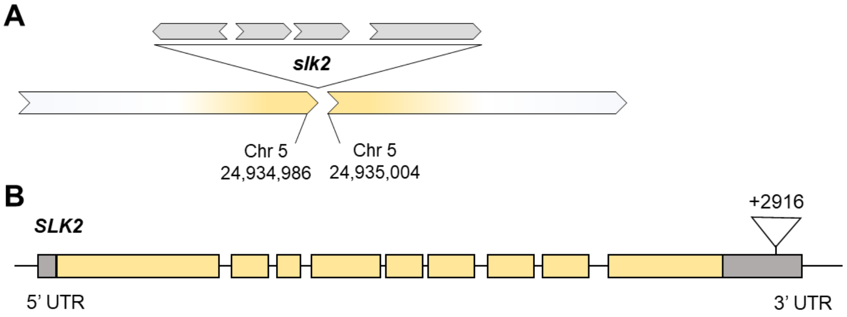


C

| Gene | Photo period | Light intensity (μE) | Harvesting point | $log_2FC$ | padj |
|---|---|---|---|---|---|
| *SLK2* (At5g62090) | SD | 60 | -1 | -0.09 | 0.80 |
| | | | +1 | -0.08 | 0.72 |

**Supplementary Figure 1. Off-target T-DNA insertion in *SLK2* and its expression in *mdh1xme1xme2.1*. A)** Long-read sequencing of *mdh1xme1xme2.1* revealed a T-DNA insertion in the 3´UTR of *SLK2* (At5g62090). Accessed by ONT sequencing. Gray blocks depict individual T-DNA copies; arrowheads indicate orientation; schematic not to scale. **B)** *SLK2* gene model indicating the location of the T-DNA insertion. Exons are represented in yellow, the UTR regions in gray, and the spaces indicate the intron regions. **C)** *SLK2* expression relative from wild-type calculated by $log_2FC$ from RNAseq (Suppl. Data 1) using DESeq2 package in R studio. Three biological replicates were used from 40-days-old plants grown in SD at LL. Tissue was harvested 1 h before (-1) and 1 h after (+1) onset of the light. Statistical significance versus wild type was assessed by Wald test using DESeq2 R package. padj, adjusted p-value.

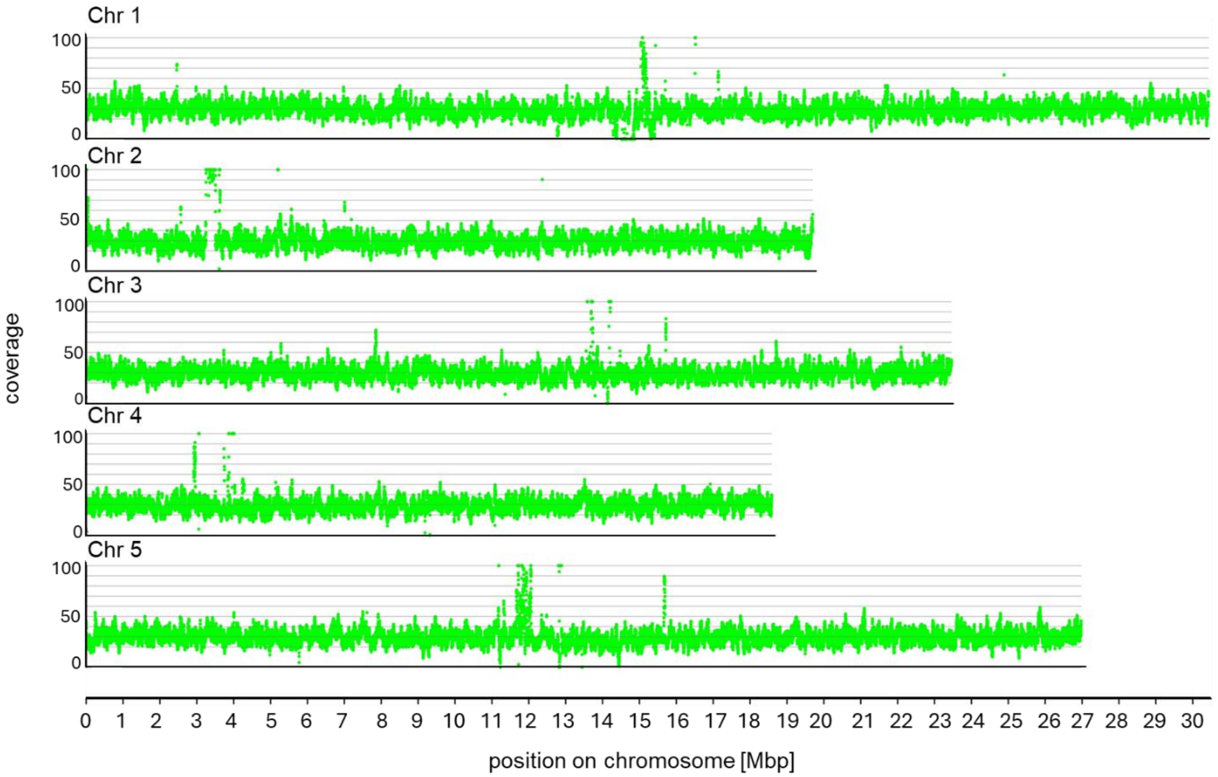


**Supplementary Figure 2. Analysis of chromosome re-arrangements of the *mdh1xme1xme2.1* nuclear genome.** Read coverage depth plot using Python scripts at Github: https://github.com/bpucker/GKseq2. Coverage peaks are observed around the centromeric regions can be attributed to incompletely and inaccurately resolved regions in the TAIR10 reference genome sequence. Chr: chromosome

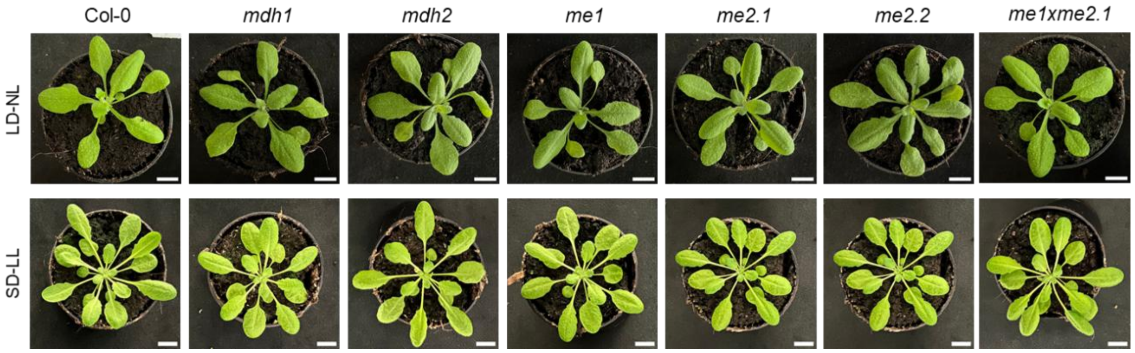


**Supplementary Figure 3. Phenotypes of different mutants in LD and SD.** Representative plants of each genotype at 21 DAS in LD at NL and 40 DAS in SD at LL. Scale bar = 1 cm. LD: long day; LL: low light; NL: normal light; SD: short day.

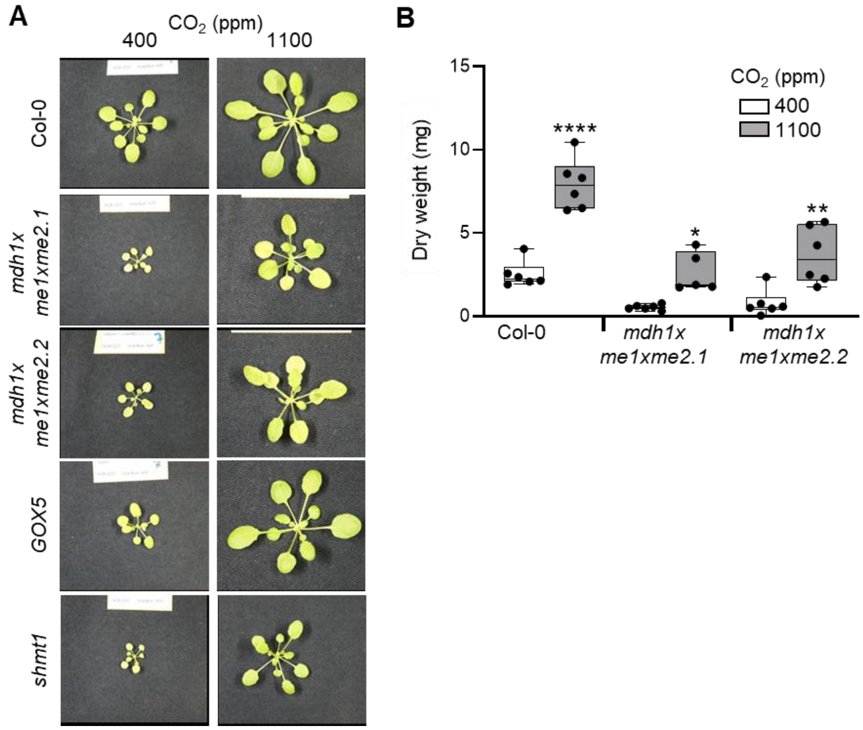


**Supplementary Figure 4. Impact of elevated $CO_2$ on leaf gas exchange and rosette biomass under LL-SD. A)** Growth of plants under normal and high $CO_2$ concentrations. Representative plants of each genotype grown during 47 days in SD conditions at 400 ppm $CO_2$ (normal) or 1100 ppm $CO_2$ (high) under LL. Photorespiratory controls (GOX5, *shmt1*) show the expected growth stimulation under elevated $CO_2$. **B)** Rosette dry weight of Col-0 and triple mutants in B. Values are means ± SD (n = 5-7). Testing for significant differences was performed by the Welch's t-test. The asterisk (*) indicates significant differences between the values under normal and high $CO_2$ conditions ($p < 0.05$, *; $p < 0.01$, **; $p < 0.001$, ***; $p < 0.0001$, ****).

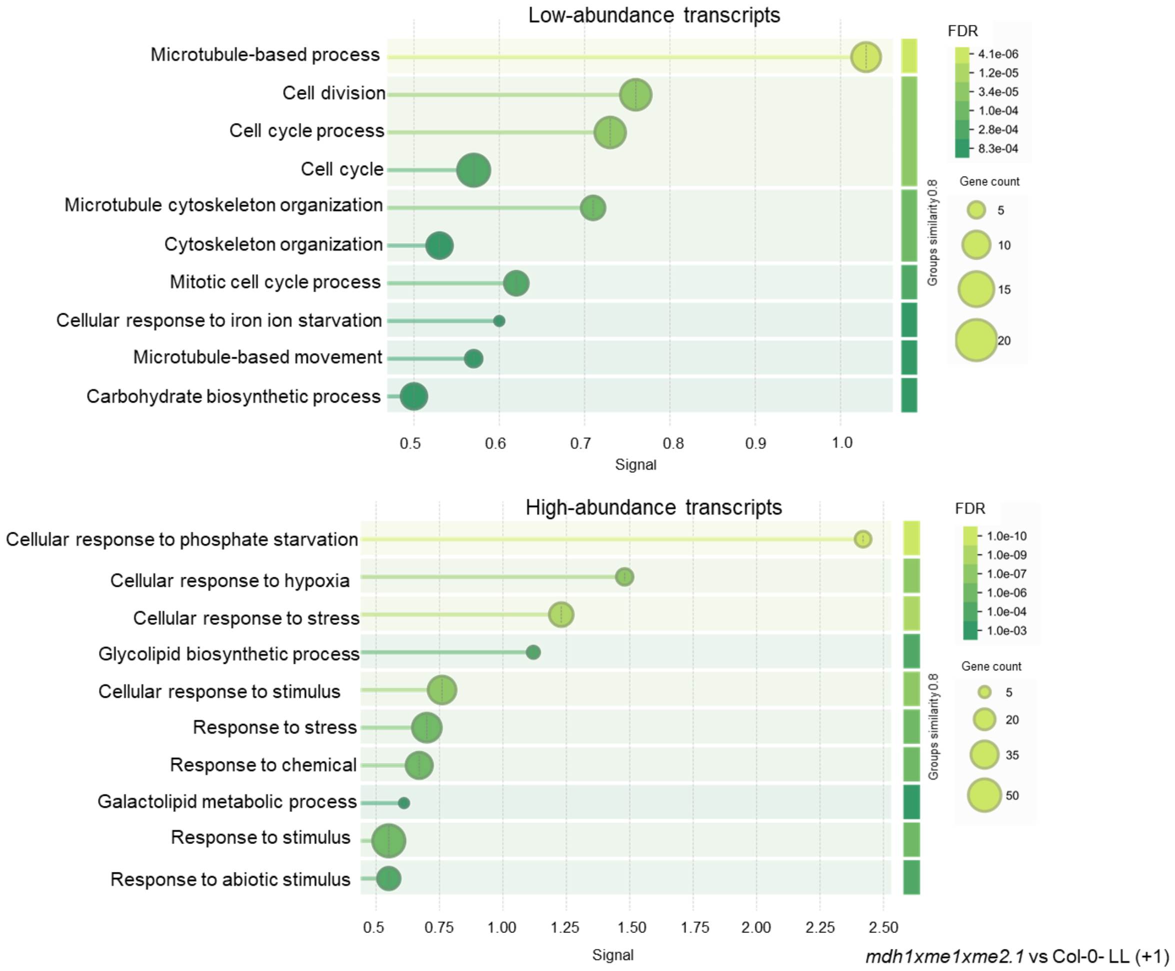


**Supplementary Figure 5. Most affected biological processes at the transcriptional level in *mdh1xme1xme2.1* under LL one hour after light turns on.** GO term enrichment analysis by STRING using the uniquely high-abundance or low-abundance transcripts at LL +1 (selected by Venn Diagram) to identify GO terms that are over-represented (or under-represented) in this condition. FDR: describes how significant the enrichment is. Shown are p-values corrected for multiple testing within each category using the Benjamini–Hochberg procedure. Signal: signal is defined as a weighted harmonic mean between the observed/expected ratio and -log(FDR). FDR tends to emphasize larger terms due to their potential for achieving lower p-values, while the observed/expected ratio highlights smaller terms, which have a high foreground to background ratio but cannot achieve low FDR values due to their size. The signal measure seeks to balance both metrics for more intuitive ordering of enriched terms. Supplemental Data 1.

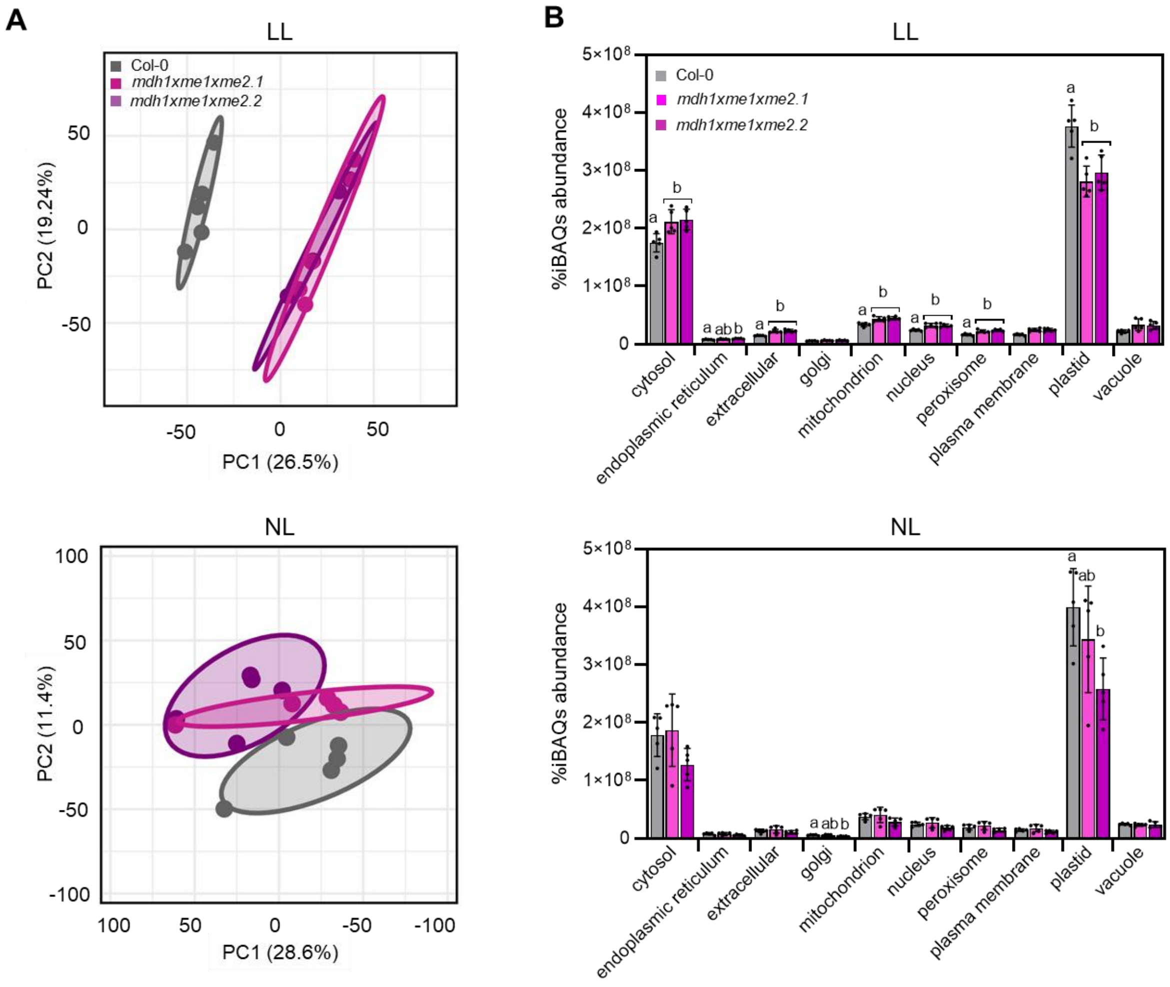


**Supplementary Figure 6. Light-dependent proteome changes between Col-0 and mutant plants under SD. A)** Principal component analysis of proteomic data obtained by mass spectrometry from plants grown at LL and NL. Confidence ellipse 80%. **B)** Relative protein abundance, expressed as percentage of total iBAQ (intensity-Based Absolute Quantification) values across subcellular compartments of plants grown at LL and NL. Data are means ± SD (n = 5). Different letters indicate significant differences ($p < 0.05$) according to ANOVA followed by Tukey's post-hoc test. LL: low light; NN: normal light; SD: short day.

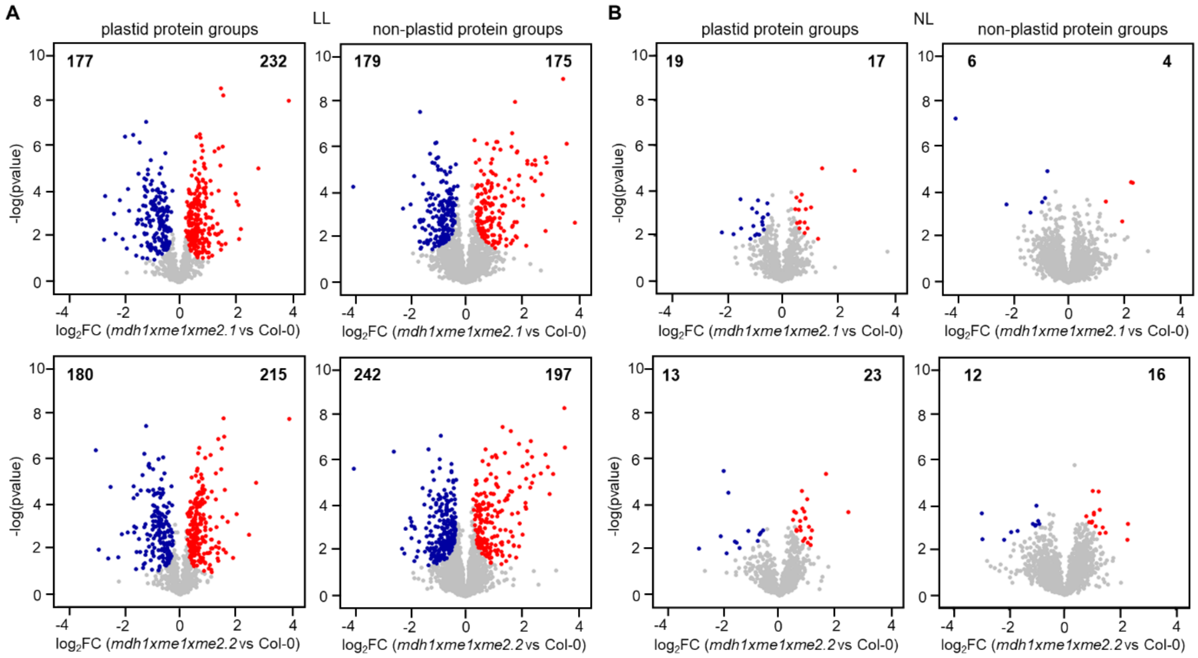


**Supplementary Figure 7. Light-dependent abundance changes in plastid and non-plastid protein groups between *mdh1xme1xme2.1/2* and Col-0 grown under LL and NL in SD.** Comparison of $log_2$-transformed LFQ-values of *mdh1xme1xme2.1/2* and Col-0 plastid and non-plastid protein groups (classified by SUBA5) grown in SD at (**A**) LL or (**B**) NL. Statistical thresholds were set at FDR $\leq 0.05$ and $S_0 \geq 0.1$. Protein groups were graphed by fold change ($log_2FC$; x-axis) and the confidence statistic ($-log_{10}$pvalue; y-axis). Blue dots represent proteins that are of significantly lower abundance in *mdh1xme1xme2.1/2* while red dots represent proteins that are significantly higher abundance in *mdh1xme1xme2.1/2* compared to Col-0. Gray dots represent the protein groups that do not differ in abundance. Numbers in the top left and right corners indicate protein groups of lower and higher abundance, respectively. LL: low light; NL: normal light; SD: short day.

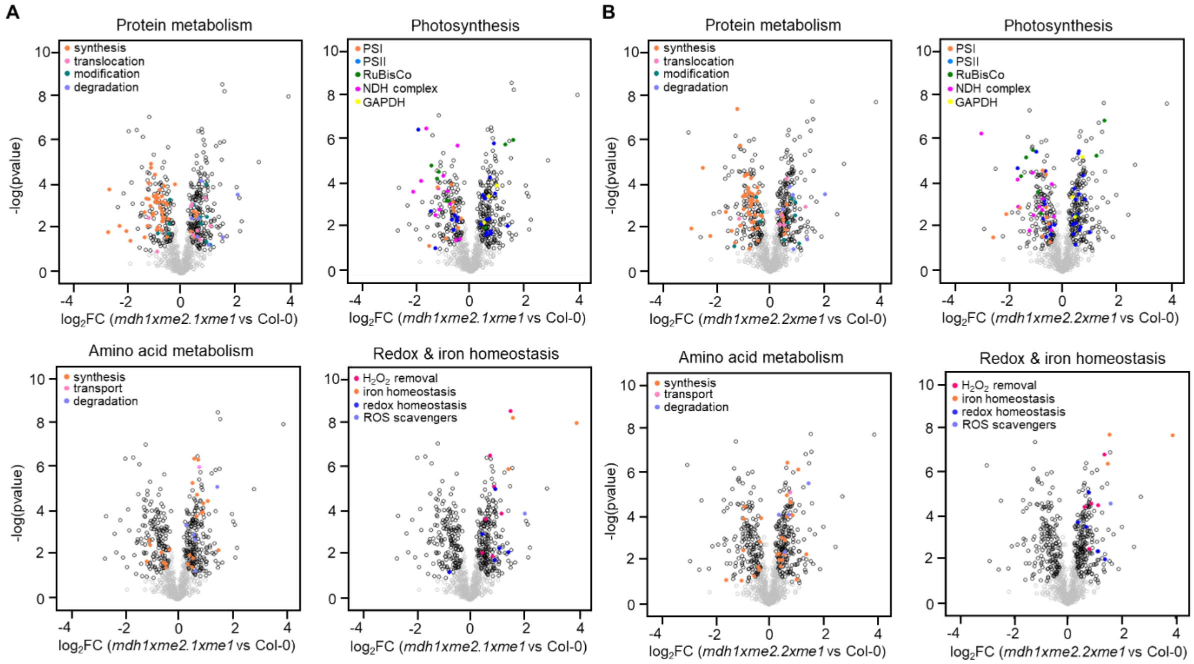


**Supplementary Figure 8. Most affected biological processes at the proteomic level in plastids from *mdh1xme1xme2.1/2* grown under LL**. After normalization, log2-transformed LFQ values of protein groups assigned to plastid by the SUBAcon algorithm were used to assess differences in the chloroplast proteome of **(A)** *mdh1xme1xme2.1* and Col-0, and **(B)** *mdh1xme1xme2.2* and Col-0 by statistical testing employing five independent biological replicates. Protein groups were classified by MapMan annotation in different levels based on their biological function and the significantly most affected biological process with their subcategories were represented in each panel with different colors (see figure legend). The complete list of categories is provided in Supplemental Data 2. Statistical thresholds were set at FDR $\leq 0.05$ and $S_0 \geq 0.1$. Black dots represent protein groups that significantly differ from *mdh1xme1xme2.1/2* to Col-0 but do not belong to the represented biological pathways, while gray dots represent protein groups below the significance (FDR) threshold. $log_2FC$, fold change; $-log_{10}$(pvalue), confidence statistics t-test. LL: low light; NN: normal light.

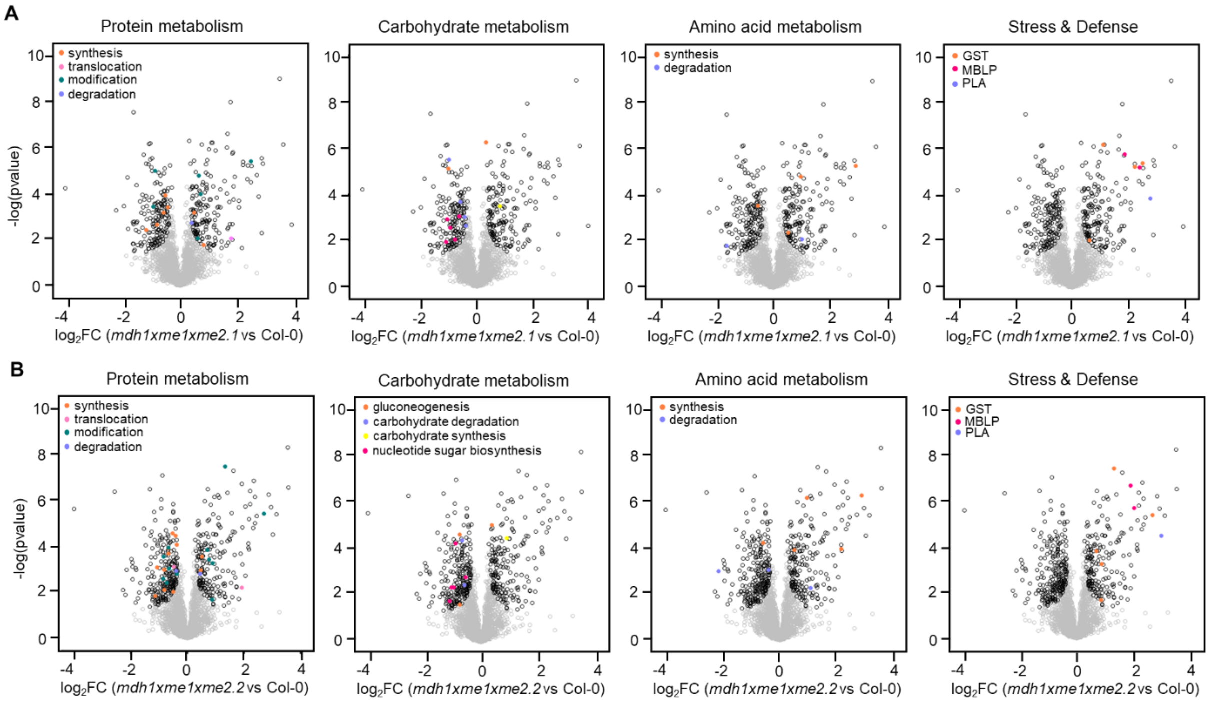


**Supplementary Figure 9. Most affected biological processes at the proteomic level in the *mdh1xme1xme2.1/2* cytosol grown under LL.** After normalization, log2-transformed LFQ values of protein groups not assigned to plastid by the SUBAcon algorithm were used to assess differences in the cytosolic proteome of **(A)** *mdh1xme1xme2.1* and Col-0 and **(B)** *mdh1xme1xme2.2* and Col-0 by statistical testing employing five independent biological replicates. Significant different abundant proteins were classified by MapMan annotation in levels based on their biological function and the most affected biological process with their subcategories were represented in each panel with different colors (see figure legend). The complete list of categories is provided in Supplemental Data 2. Statistical thresholds were set at FDR ≤ 0.05 and $S_0 \geq 0.1$. Black dots represent protein groups that significantly differ from *mdh1xme1xme2.1/2* to Col-0 but do not belong to the represented biological pathways, while gray dots represent protein groups below the significance (FDR) threshold. $\log_2$FC, fold change; $-\log_{10}$(pvalue), confidence statistics T-test. GST: glutathione S-transferases; MBLP: mannose-binding lectin proteins; PLA: phospholipase A; LL: low light.

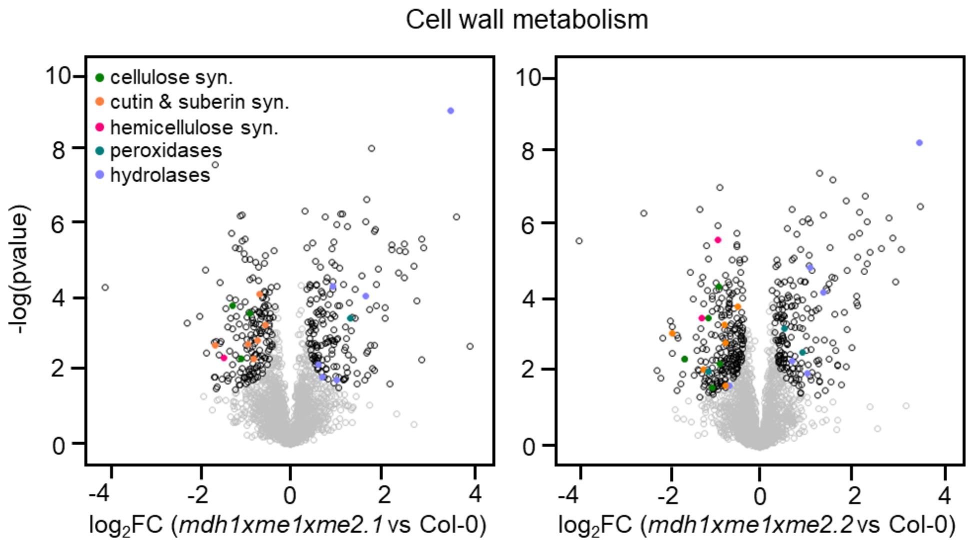


**Supplementary Figure 10. Protein groups abundance changes cell wall metabolism in *mdh1xme1xme2* under LL.** After normalization, log2-transformed LFQ values of protein groups not assigned to plastid by the SUBAcon algorithm were used to assess differences in the proteome between *mdh1xme1xme2.1* and Col-0, and *mdh1xme1xme2.2* and Col-0 by statistical testing employing five independent biological replicates. Protein groups were classified by MapMan annotation in levels based on their biological function. The complete list of categories is provided in Supplemental Data 2. Statistical thresholds were set at FDR < 0.05 and $S_0 = 0.1$. Black dots represent protein groups that significantly differ from *mdh1xme1xme2* to Col-0 but do not belong to the represented biological pathway, while gray dots represent protein groups lower the significant threshold. LL: low light; syn: synthesis.

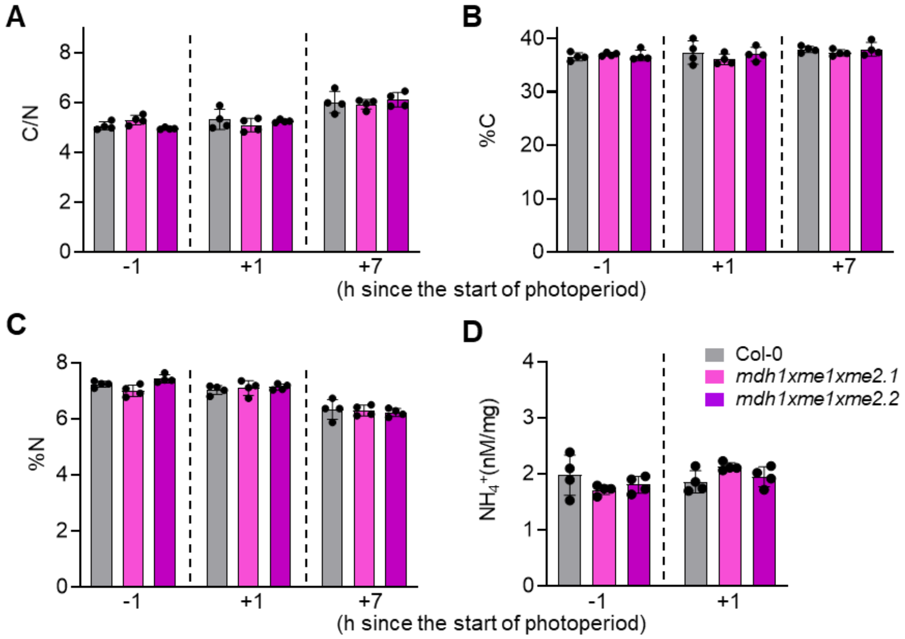


**Supplementary Figure 11. C/N ratio, total C, total N and ammonia in plants grown in SD and NL. A)** Carbon-to-nitrogen ratio (C/N ratio), calculated as total carbon divided by total nitrogen. **B)** C content (%C) and **C)** N content (%N) expressed as percentage of dry weight in samples harvested 1 h before light onset (-1), one hour after light onset (+1), and one hour before light offset (+7). **D)** Leaf total ammonia ($NH_4^+$) concentrations measured one hour before light onset (-1) and one hour after light onset (+1). Data represent means ± SD of 3-4 biological replicates per line. Different letters indicate significant differences ($p$ < 0.05) according to ANOVA followed by Tukey's post-hoc test. NL: low light; SD: short day.

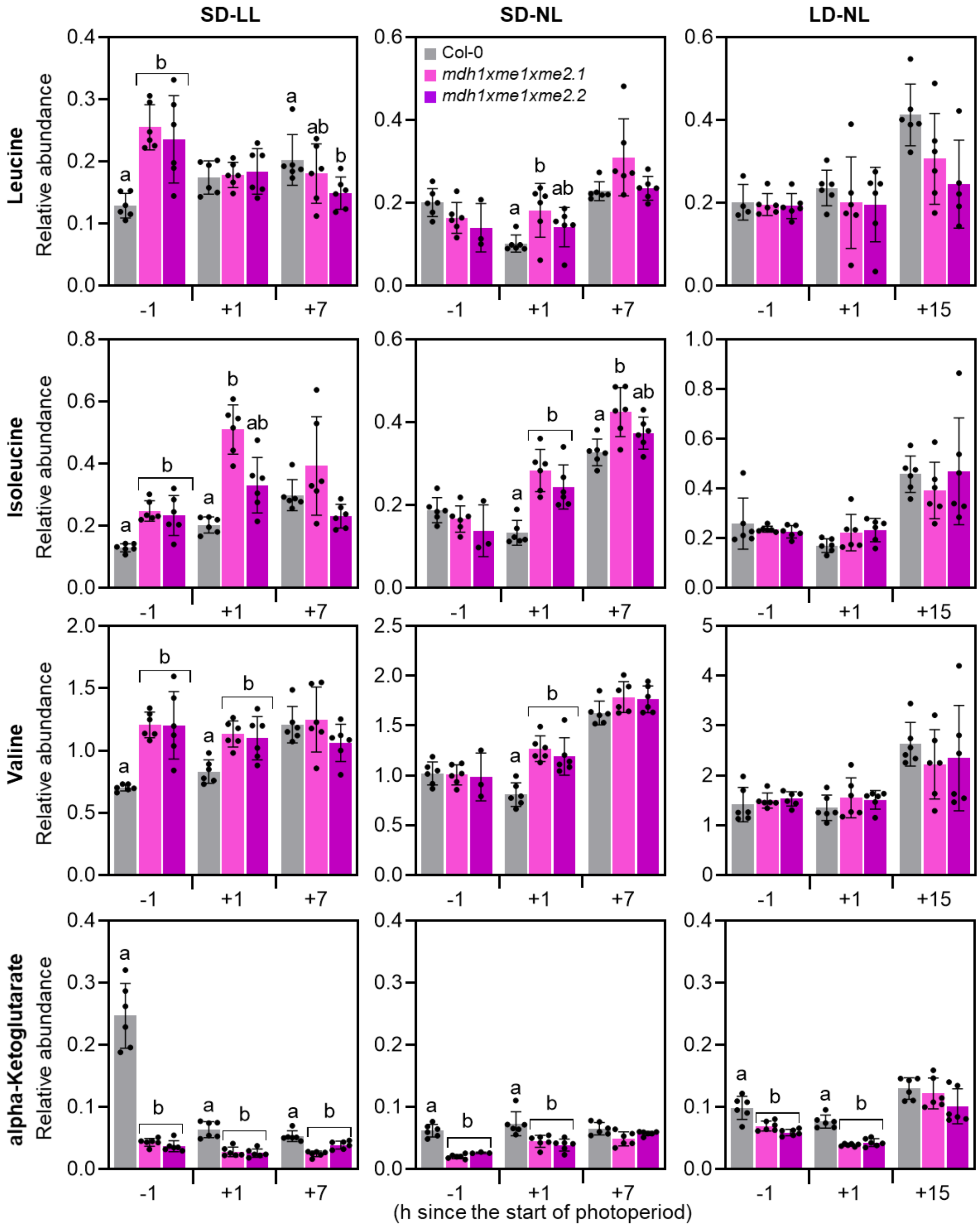


**Supplementary Figure 12. Relative concentrations of BCAA and alpha-Ketoglutarate.** Relative levels were quantified by GC-MS in leaves of the indicated genotypes grown under LL SD, NL SD and NL LD. Samples were harvested 1 h before lights-on (-1 h), 1 h after lights-on (+1 h) and 1 h before lights-off (+7/+15 h). Bars show means ± SD (n = 6 independent biological replicates). Statistics were performed separately for each time point using one-way ANOVA followed by Tukey’s post-hoc test; bars that do not share a letter differ significantly at $p < 0.05$. LD: long day; LL: low light; NL: normal light; SD: short day.

#### Supplementary Tables (Martinez et al., 2026)

**Supplementary Table 1.** Information on the T-DNA insertion lines used in the work. s. a. = see above; ko = knock out; kd = knock down.

| Mutant (Type) | Gene | Gene locus | Insertion line | Reference | Phenotype in LL SD |
|---|---|---|---|---|---|
| *me1* (ko) | ME1 | At2g13560 | Sail-374-A02 | Tronconi et al., 2008 | no |
| *me2.1* (ko) | ME2 | At4g00570 | Sail-291-C05 | | no |
| *me2.2* (kd) | ME2 | At4g00570 | SALK_131720 | | no |
| *me1xme2.1* | s. a. | s. a. | s. a. | | no |
| *mdh1* (ko) | MDH1 | At1g53240 | GABI_097C10 | Tomaz et al., 2010 | no |
| *mdh1xme1xme2.1* | s. a. | s. a. | s. a. | This work | yes |
| *mdh1xme1xme2.2* | s. a. | s. a. | s. a. | | yes |

**Supplementary Table 2.** List of primers used in this study. s. a. = see above

| **Name** | **Sequence (5'- 3')** | **Gene/T-DNA Insertion** | **AGIs** |
|---|---|---|---|
| me1-Fw1 | ACGATGACGGAGAGAAATCGT | *ME1* | At2g13560 |
| me1-Rv1 | CTGGACATCATCATTGAACAT | | |
| me2-Fw1 | GACCTGTGTACAGCAATGTGATCG | *ME2* | At4g00570 |
| me2-Rv1 | CCTGGACATCATCATTGAACATGC | | |
| mdh1-Fw1 | TCCGATCTTCTGCCTCC | *MDH1* | At1g53240 |
| mdh1-Rv1 | CAACTGGGACATTTGCCTT | | |
| act2-Fw1 | TAACTCTCCCGCTATGTATGTCGC | *ACT2* | At3g18780 |
| act2-Rv1 | GAAGCAAGAATGGAACCACCG | | |
| Sail | TAGCATCTGAATTTCATAACCAATCTCGA TACAC | *ME1 / ME2.1* | s. a. |
| SALK | TGGTTCACGTAGTGGGCCATCG | *ME2.2* | s. a. |
| GABI | ATATTGACCATCATACTCATTGC | *MDH1* | s. a. |
| me1-Fw2 | AAGATGAAGATCGTTGTTGCTG | *ME1* | s. a. |
| me1-Rv2 | CATCAACCACCCAAAATTGACT | | |
| me2-Fw2 | TTGGTGTAACTAAGATGGCAGT | *ME2* | s. a. |
| me2-Rv2 | CGTATCTCAGCTGGATTTTTGG | | |
| mdh1-Fw2 | TGTCCTCTCCGTGGTGACAAGACC | *MDH1* | s. a. |
| mdh1-Rv2 | TCCGATCTTCTGCCTCC | | |
| mdh2-Fw2 | CGGTCACATCAACACCCGAT | *MDH2* | s. a. |
| mdh2-Rv2 | AGCGCCTTCTAGAGCTTTCC | | |
| act2-Fw2 | CTTGCACCAAGCAGCATGAA | *ACT2* | s. a. |
| act2-Rv2 | CCGATCCAGACACTGTACTTCCTT | | |
